# Systematic Engineering of Artificial Metalloenzymes for New-to-Nature Reactions

**DOI:** 10.1101/2020.07.15.204206

**Authors:** Tobias Vornholt, Fadri Christoffel, Michela M. Pellizzoni, Sven Panke, Thomas R. Ward, Markus Jeschek

## Abstract

Artificial metalloenzymes (ArMs) catalyzing new-to-nature reactions under mild conditions could play an important role in the transition to a sustainable, circular economy. While ArMs have been created for a variety of bioorthogonal transformations, attempts at optimizing their performance by enzyme engineering have been case-specific and resulted only in modest improvements. To realize the full potential of ArMs, methods that enable the rapid discovery of highly active ArM variants for any reaction of interest are required. Here, we introduce a broadly applicable, automation-compatible ArM engineering platform, which relies on periplasmic compartmentalization in *Escherichia coli* to rapidly and reliably identify improved ArM variants based on the biotin-streptavidin technology. We systematically assess 400 ArM mutants for five bioorthogonal transformations involving different metal cofactors, reaction mechanisms and substrate-product pairs, including novel ArMs for gold-catalyzed hydroamination and hydroarylation. The achieved activity enhancements of up to fifteen-fold over wild type highlight the potential of the systematic approach to ArM engineering. We further capitalize on the sequence-activity data to suggest and validate smart strategies for future screening campaigns. This systematic, multi-reaction study has important implications for the development of highly active ArMs for novel applications in biocatalysis and synthetic biology.

## Introduction

Artificial metalloenzymes (ArMs) combine the broad reaction scope of organometallic catalysis with the exceptional catalytic performance, selectivity and mild reaction conditions of enzymes^1,2^. Therefore, they have a great potential to enable sustainable synthetic routes to various compounds of interest^3,4^ and, if functional in a cellular environment, open up new possibilities for metabolic engineering and synthetic biology^5^. ArMs have been constructed by repurposing natural metalloenzymes^6,7^, designing binding sites for metal ions^8–11^ and incorporating organometallic cofactors into protein scaffolds^12–20^. Moreover, unnatural amino acids^21^ and photoexitation^22,23^ have been used to unlock new reactivity in enzymes. Amongst these efforts to create new-to-nature biocatalysts, one of the most versatile approaches relies on the biotin-streptavidin technology. This strategy employs the high affinity of the homotetrameric protein streptavidin (Sav) for the vitamin D-biotin to non-covalently anchor biotinylated metal complexes (referred to as cofactor hereafter) within the Sav protein. Using this approach, artificial enzymes have been created for multiple reactions, including hydrogenation, sulfoxidation, C−H activation and olefin metathesis^24–26^. While in some cases wild-type Sav imparts some selectivity or rate acceleration on the reaction, protein engineering is usually required in order to obtain proficient biocatalysts^27–31^. Initial studies in this direction relied on screening (semi-)purified Sav mutants^32,33^, but more recently a trend towards whole-cell screening with *Escherichia coli* has emerged for Sav-based ArMs^34,35^ as well as other artificial enzymes^8,36,37^. This allows significantly increased throughput and is the method of choice if variants that are functional under *in vivo* conditions are desired, which is an important prerequisite for synthetic biology applications.

However, a number of challenges arise for cell-based ArM assays, most notably insufficient cofactor uptake into the cell as well as cofactor poisoning by cellular components such as reduced glutathione^38^. In order to circumvent these challenges, Sav has been exported to the periplasmic space^34^ or the cell surface^35^. While these studies have established the possibility of engineering ArMs using whole-cell screenings, they did so using case-specific engineering strategies and with a focus on reactions affording fluorescent products. Consequently, general applicability of these methods across various reactions remains to be demonstrated. To generalize the development of ArMs, broadly applicable engineering strategies that enable the rapid identification of highly active variants for ideally any reaction of interest are needed. This requires a robust screening protocol that is compatible with various reaction conditions and analytical readouts. Moreover, it imposes a demand for Sav mutant libraries that embody a high potential to contain highly active variants for different reactions while maintaining a comparably small and thus screenable library size. Such a combination of a broadly applicable screening method with a “concise” library could serve as a universal starting point for various ArM engineering campaigns and would render tedious case-by-case method and library development obsolete. Herein, we present a screening platform that meets the aforementioned requirements as a first systematic approach to ArM engineering. We establish a well-plate based screening protocol that can be easily adapted to new reaction conditions or analytical methods and show that it is amenable to lab automation as an important prerequisite for streamlined ArM engineering. Further, we create a full-factorial, sequence-defined Sav library that is rich in variants with high activity by simultaneously diversifying two crucial amino acid positions in close vicinity to the catalytic metal. To demonstrate the versatility of this platform, we selected five bioorthogonal reactions requiring different cofactors, reaction conditions and analytical readouts. The platform enabled the identification of substantially improved ArMs for all tested reactions with fast turnaround times. Moreover, the systematic characterization of the local sequence-activity landscape enabled us to identify smart library designs to further enhance the development of ArMs. This study represents the first systematic approach to ArM engineering and thus constitutes an important step towards a streamlined optimization of these novel biocatalysts.

## Results

### Construction and characterization of a sequence-defined Sav library

In order to establish a generalizable first engineering step for ArMs, we sought to generate a library of Sav variants that offers a high likelihood of identifying active variants at a minimized screening effort. Based on previous experience with Sav expression and whole-cell ArM catalysis^34,35^, we selected a periplasmic compartmentalization strategy for this library. Secretion to the periplasm and subsequent ArM assembly represents a good trade-off between accessibility to the cofactor, expression levels and compatibility of the reaction conditions^39^. In order to facilitate periplasmic export in *E. coli*, the signal peptide of the outer membrane protein A (OmpA) was N-terminally fused to T7-tagged mature Sav (referred to as wild type hereafter) as previously reported^34^. The holoenzyme (i.e. full ArM with cofactor) can then be assembled in the periplasm by incubating cells in a buffer containing a biotinylated metal cofactor (Fig. 1a). We selected the amino acid residues 112 and 121 in Sav as randomization targets, which correspond to serine and lysine in wild-type Sav. These residues are in close proximity to the biotinylated cofactor^32^ (Fig. 1b) and have repeatedly been found to have a substantial impact on the activity of ArMs for diverse cofactors and reactions^33–35^. More specifically, we created a combinatorial Sav 112X 121X library (X representing all 20 canonical amino acids) consisting of all 400 possible amino acid combinations for these two positions. Such a full-factorial library makes it possible to identify improved mutants that are the result of synergistic interactions between the two positions. We applied a three-step cloning and sequencing strategy (see Methods and Supplementary Figs. 1-3) to produce an arrayed, sequence-verified set of all 400 possible Sav 112X 121X mutants. This minimizes the screening effort by avoiding the requirement for oversampling.

**Figure 1.**
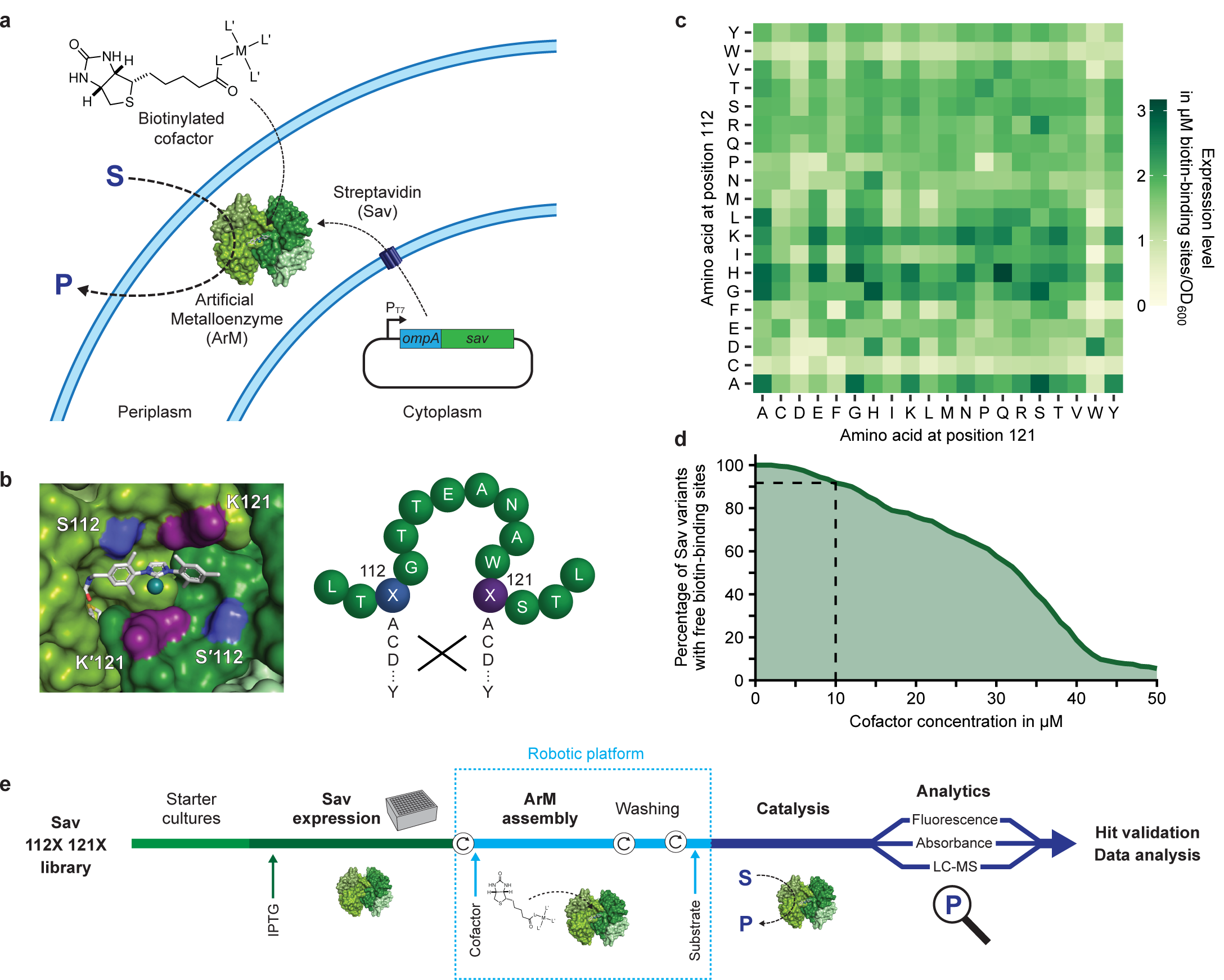
Platform for the systematic screening of ArMs in *E. coli*’s periplasm. **a**, By fusion to the signal peptide of the outer membrane protein A (OmpA), Sav is secreted to the periplasm, where it binds to an externally-added biotinylated cofactor to afford an ArM. The cofactor consists of a catalytically active metal M and its ligands L and L’. **b**, Left: Close-up view of a bound biotinylated metathesis cofactor within the biotin-binding vestibule of homotetrameric Sav (PDB 5IRA). The symmetry-related amino acids S112 & S’112 (blue) and K121 and K’121 (purple) in Sav are located in the immediate vicinity of the cofactor and were mutated in a combinatorial manner (right), affording an Sav library with 400 amino acid combinations in the two positions. **c**, Expression level of the 400 Sav mutants as determined in cell lysate using a fluorescence quenching assay40 (Methods). The values displayed are the mean of biological triplicates (n=3) and were normalized to the OD600 of the cultures (refer to Supplementary Figure 4 for the standard deviation between replicates). **d**, Estimated percentage of Sav mutants with unoccupied biotin-binding sites as a function of the concentration of cofactor added to the cell suspension (assuming 50 % uptake into the periplasm). 10 µM were selected as the maximum permitted cofactor concentration for further experiments since this leads to a maintained excess of binding sites for more than 90 % of Sav variants (dashed lines). **e**, Overview of the screening workflow (Methods). Following periplasmic Sav expression in deep-well plates, cells are incubated with the respective biotinylated cofactor and subsequently washed once prior to the addition of substrate (circular arrows represent centrifugation steps for buffer exchange). Depending on the reaction, the product concentration is determined via absorbance or fluorescence measurements or by LC-MS.

Even single mutations can substantially alter the expression level of proteins^41^, which complicates the identification of variants with increased specific activity. For this reason, we aimed to screen at cofactor concentrations that do not fully saturate the available biotin-binding sites even for low-expressing variants (i.e. excess of binding sites). Consequently, we determined the expression level of all 400 Sav mutants from the 112X 121X library, relying on a quenching assay with biotinylated fluorescent probes^40^ (Fig. 1c, Methods). The majority of variants showed high expression levels ranging from 17 to 536 mg L^-1^, which is equivalent to concentrations of 1 to 33 µM of biotin-binding sites. When normalized by the density of the cell suspensions, these values amount to 0.2 to 3.2 µM binding sites in a culture with an optical density at 600 nm (OD_600_) of one. Importantly, while the expression of variants harboring a cysteine at position 112 or a tryptophan at either of the two positions appeared to be reduced, 87 % of Sav mutants showed an expression level greater than 1 µM binding sites per OD_600_. Based on these measurements, we determined a maximum permitted cofactor concentration (Fig. 1d). This critical experimental parameter should on the one hand be high enough to result in well-detectable product concentrations and on the other hand remain below the concentration of available biotin-binding sites for the majority of the library. The latter requirement is important in order to keep the concentration of assembled ArMs constant, irrespective of the expression level of Sav. In light of previous studies^34^ and our own observations (Supplementary Fig. 5), we conservatively assumed that less than half of externally-added cofactor enters the periplasm. As a consequence, we set the maximum permitted cofactor concentration at the incubation step to 10 µM because this ensures that an excess of binding sites is maintained for more than 90 % of variants (Fig. 1d), thus largely eliminating the expression level dependence of the screening results.

### Systematic screening of ArMs for bioorthogonal reactions

In order to facilitate screening of improved ArMs for diverse reactions, we relied on a 96-well-plate based assay^34^. In brief, periplasmic expression cultures are spun down and cells are resuspended in an incubation buffer containing the cofactor of interest to assemble the ArM in the periplasm. After incubation, cells are washed once to remove unbound cofactor and resuspended in a reaction-specific buffer containing the substrate. Following overnight incubation, the product concentration is determined using a suitable analytical method (Fig. 1e). This screening procedure is compatible with various reaction conditions and analytical methods, and consequently not restricted to model reactions, which, for example, produce a fluorescent product. Furthermore, it is amenable to lab automation (see below). Relying on this protocol, we sought to systematically test the full-factorial 400-mutant Sav library for various ArM reactions of interest (Fig. 2a). To this end, we selected three reactions based on previously reported ArMs: A ring-closing metathesis (RCM, **I**) of diallyl-sulfonamide **1** to 2,5-dihydro-pyrrole **2**^34^ and two deallylation reactions (**II, III**) of allyl carbamate-protected substrates **3** and **5**, affording amino coumarin **4** and indole **6**, respectively. The corresponding cofactors for these reactions are a biotinylated second-generation Hoveyda-Grubbs catalyst **(Biot-NHC)Ru** (reaction **I**) and a biotinylated ruthenium cyclopentadienyl complex **(Biot-HQ)CpRu**^42,43^ (reactions **II** and **III**), respectively (Fig. 2b). Further, with the aim of extending the scope of ArM-catalyzed reactions towards biocompatible nucleophilic cyclizations of alkynes, we developed two novel gold-containing ArM cofactors, (**Biot-NHC)Au1** and (**Biot-NHC’)Au2** (Fig. 2b), to create ArMs for hydroamination (**IV**) and hydroarylation^44^ (**V**) reactions. Importantly, none of these reactions are known to be catalyzed by natural enzymes, and therefore they have potential for applications ranging from the design of novel metabolic pathways to prodrug activation and *in vivo* labelling^5,45^.

**Figure 2.**
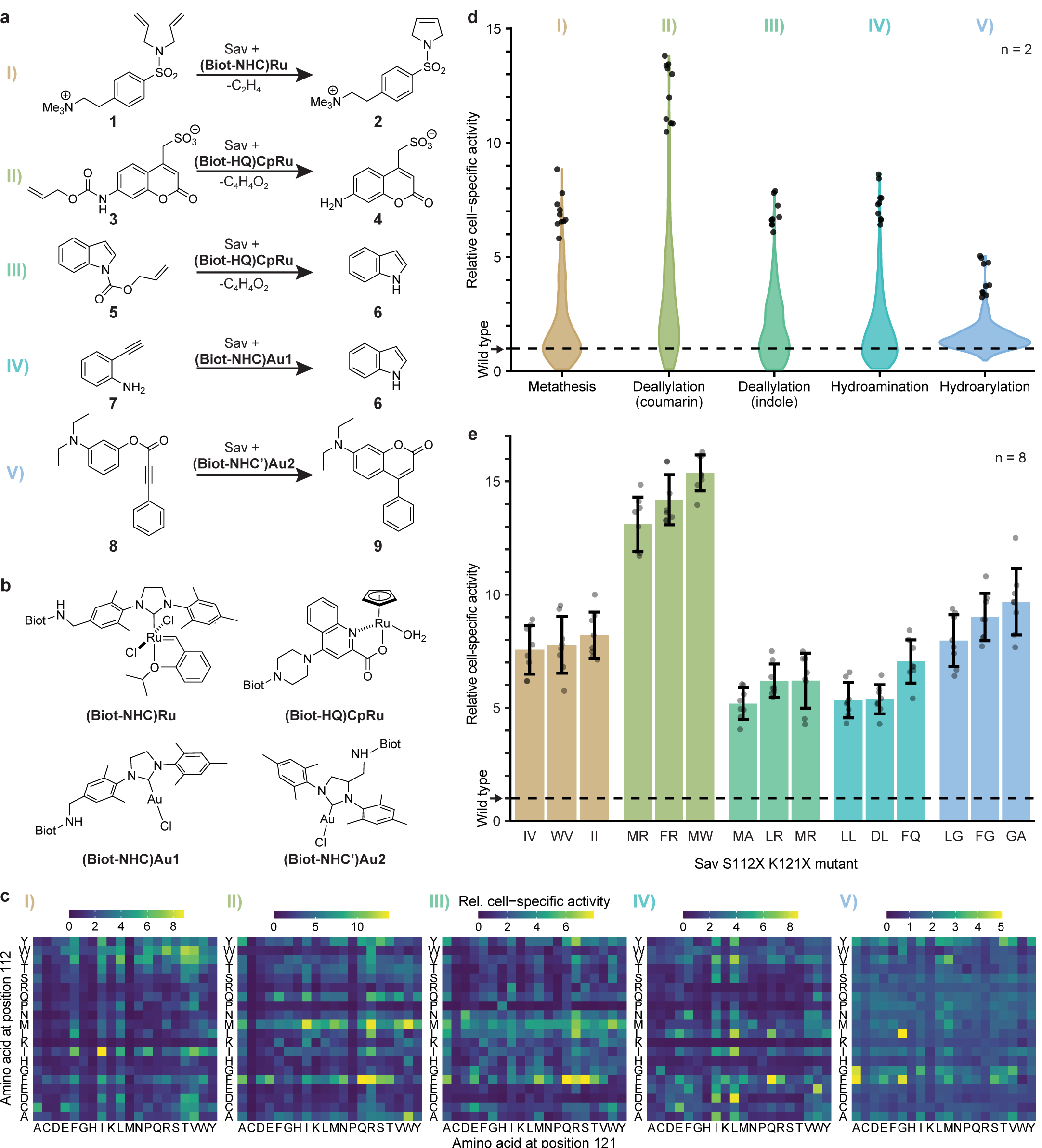
Systematic screening for ArMs catalyzing diverse reactions. **a**, ArM-catalyzed reactions: **I**, Ring-closing metathesis (RCM) with a diallyl-sulfonamide **1** to afford a 2,5-dihydro-pyrrole **2. II**, Deallylation of allyl carbamate-protected coumarin **3** to the corresponding amino coumarin **4. III**, Deallylation of allyl carbamate indole **5** to indole **6. IV**, Hydroamination of 2-ethynylaniline **7** to indole **6. V**, Hydroarylation of profluorophore **8** to afford amino coumarin **9. b**, Structures of the biotinylated cofactors employed in this study. Biot refers to D-biotin as depicted in Figure 1a. **c**, Cell-specific activity of 400 ArMs mutated at positions 112 and 121 in Sav normalized to the activity of wild-type Sav (S112 K121). The displayed activities correspond to the product concentrations after 20 h of reaction and are the mean of biological duplicate reactions. The corresponding standard deviations between replicates can be found in Supplementary Figure 8. Note that the screenings for reactions **II** and **V** were performed using the robotic platform. **d**, Activity distribution in the Sav mutant library for the five different ArM reactions. Violins comprise all 400 double mutants with the ten most active ArMs depicted as circles. **e**, Validation of “hits” from the 400-mutant screens. Bars correspond to the mean activity of eight biological replicates with standard deviation shown as error bars and individual replicates as circles. Mutants are designated by the amino acids in position 112 and 121.

While preliminary experiments on the RCM **I** and deallylations **II** and **III** confirmed the compatibility of these reactions with the periplasmic screening platform, reactions **IV** and **V** showed only little conversion in the whole-cell assay. Based on previous experience with ArMs^34,38^, we hypothesized that cellular thiols might inhibit these gold-catalyzed reactions, which we could confirm by *in vitro* experiments with purified Sav. These revealed a marked inhibitory effect of glutathione and cysteine (Supplementary Fig. 6a). Previously, we had reported that thiol inhibition can be overcome *in vitro* by adding the oxidizing agent diamide^38^. Gratifyingly, we observed that diamide also neutralizes the detrimental effect of thiols in whole-cell experiments, rendering periplasmic gold catalysis feasible (Supplementary Fig. 6b).

With a functional periplasmic screening assay for the five reactions at hand, we tested all 400 mutants for each reaction at least in biological duplicates relying on the aforementioned workflow in 96-deep well plates. Importantly, we automated the steps required for incubation with cofactor, washing and substrate addition and applied this automated protocol for reactions **II** and **V**. This substantially reduces the manual labor and the consumption of consumables. The robotic platform can handle up to eight 96-well plates concurrently. Accordingly, the 400-member library can be processed in one day and an entire screening (including Sav expression and analytics) can be performed within one week.

Relying on the periplasmic assay, we recorded a local sequence-activity landscape for each ArM (Fig. 2c). The activity patterns varied substantially between reactions, which points to the existence of specific interactions between protein, cofactor and substrate, as opposed to unspecific effects, for instance as a result of varying expression levels. In line with this, we observed no correlation between expression level and activity for any of the reactions (Supplementary Fig. 7). Hence, the concentration of cofactor, and not the number of available biotin-binding sites, is limiting, which confirms the validity of the determined maximum cofactor concentration (see above). Similar activity patterns were only found between the two deallylation reactions (**II** and **III**), suggesting that in these cases the substrates are sufficiently similar or that the observed activity is mainly the result of interactions between protein and cofactor.

For all reactions, the wild-type variant (S112 K121) had a comparably low activity that was only slightly above the background observed for cells lacking Sav. Note that this background activity results from residual cofactor that is unspecifically bound and thus incompletely removed in the washing step. Importantly, compared to wild-type Sav we identified significantly more active mutants for all five ArM reactions (Fig. 2d). In order to validate these results, we measured the activity of the most promising variants of each ArM again in eight replicates, which confirmed the observed enhancements (Fig. 2e). The best mutants reached fold-improvements over the wild type varying between six- and fifteen-fold. Notably, we also identified substantially improved variants of the novel, gold-based ArMs for hydroamination (**IV**) and hydroarylation (**V**), reaching fold-improvements over wild type of about seven- and ten-fold, respectively.

While the screenings were designed towards increased ArM activity in a cellular environment, we sought to investigate whether the identified variants also display increased activity *in vitro*. We therefore purified the three most active mutants for each reaction and performed ArM reactions with these (Fig. 3). Notably, all mutants significantly outperformed the wild-type ArM as well as the corresponding free cofactor, as reflected by markedly higher turnover numbers (TONs). Depending on the reaction, the best variants reached between five- and twenty-fold higher TONs than the free cofactor, demonstrating the benefit of embedding these cofactors within an engineered protein scaffold. It should be noted that in some cases the relative ranking of variants changed compared to the whole-cell biotransformations, which is likely due to the different reaction conditions and the absence of cellular components *in vitro*. Nevertheless, the *in vitro* experiments confirmed that the identified variants indeed have a significantly increased specific activity and further corroborated the potential of the periplasmic screening platform for the rapid and straightforward discovery of different ArMs with improved catalytic properties.

**Figure 3.**
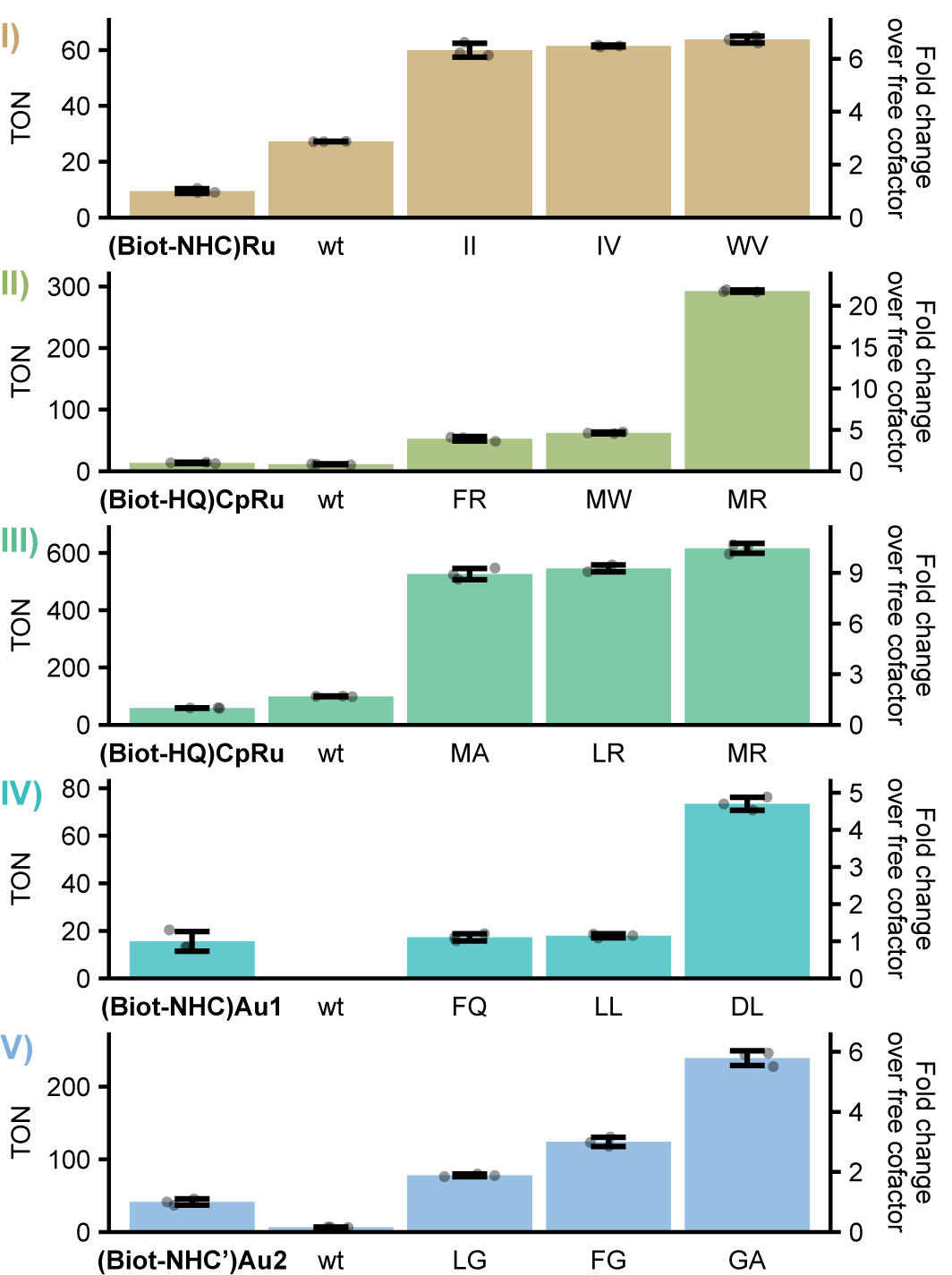
*In vitro* turnover number of ArM variants identified in the periplasmic screening. For each reaction, the three most active variants identified in the whole-cell screening were purified and their TON was determined. Reactions were carried out at 37 °C and 200 rpm for 20 h. Bars represent mean TONs of technical triplicate reactions with standard deviation as error bars and individual replicates as circles. For comparison, the free cofactor and wild-type Sav variant (wt) were included. Mutants are designated by the amino acids in position 112 and 121.

### Analysis of sequence-activity landscapes

The recording of sequence-activity data as performed herein provides the basis for additional insight beyond the mere identification of active mutants. To this end, we first sought to understand which biophysical properties result in the observed activity changes. Therefore, we analyzed the effect of individual amino acids on the activity of the different ArMs (Fig. 4a): First, we scaled the activity measurements (Fig. 2c) such that the average activity across the 400 mutant library equals zero and the corresponding standard deviation equals one. Next, we averaged the scaled activities of all 20 variants that carry the same amino acid in one position. This was performed for all 20 canonical amino acids and either of the two diversified positions. As a result, amino acid- and position-specific averages greater or smaller than zero represent a positive or negative effect on ArM catalysis, respectively. While effects of amino acids varied between positions and reactions (Fig. 4a), an overall positive effect of hydrophobic amino acids on the activity of the corresponding ArMs could be observed, whereas charged amino acids as well as proline tend towards a negative impact. Additionally, we used hierarchical clustering to analyze which amino acids have similar effects on activity. In line with the previous observation, hydrophobic amino acids clustered clearly separated from hydrophilic ones, indicating that this property is crucial in terms of catalytic activity (Fig. 4b). Further, amino acids with similar chemical properties showed a strong tendency to cluster closely together (note for instance L-I-V, D-E, H-K and S-T clusters). This points to an anticipated large potential of library strategies that employ reduced amino acid alphabets to maintain a large chemical diversity while significantly reducing the screening effort (see below).

**Figure 4.**
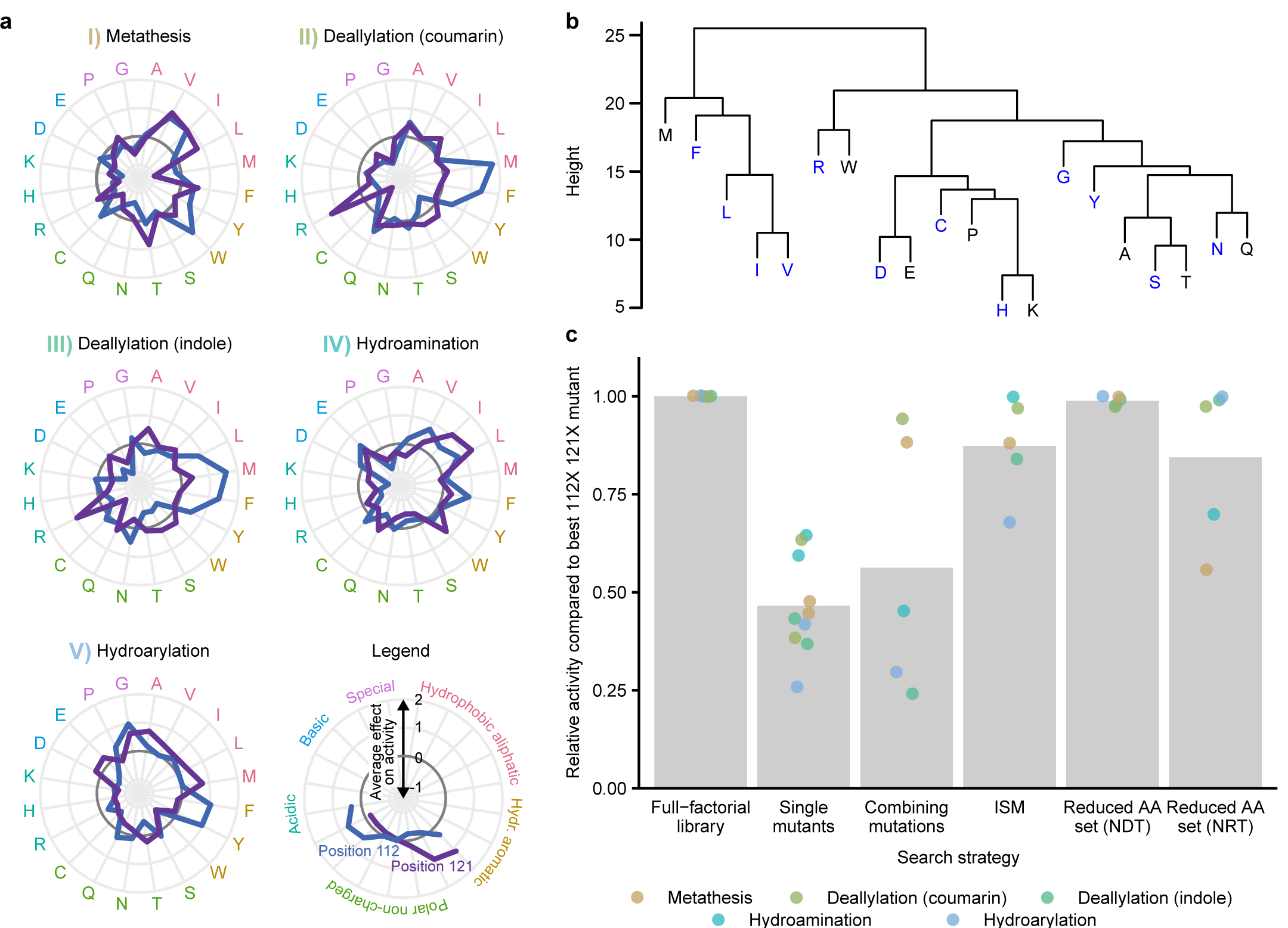
Analysis of amino acid effects and implications for directed evolution. **a**, Average effect of amino acids on ArM activity grouped by reaction and position. Data points outside the dark grey circle indicate a positive mean effect on activity, whereas data points inside denote a negative effect. Values were standardized by subtracting the mean relative activity of all 400 mutants for the respective reaction and dividing by the corresponding standard deviation. The mean across all 20 variants harboring the respective amino acid at the indicated position is shown. **b**, Hierarchical clustering of amino acids based on the activity of variants containing these residues at position 112 or 121 across all reactions (Methods). Amino acids resulting in similar activity patterns are grouped at lower heights than more dissimilar ones. Amino acids that are encoded by the degenerate codon NDT are highlighted in blue. **c**, Comparison of enzyme engineering strategies based on the results presented in this study. The success of various strategies was compared to the full-factorial approach applied herein (from left to right): Screening single mutants for both positions individually, combining the best mutations at the two individual positions, iterative saturation mutagenesis (ISM), as well as screening combinatorial libraries based on reduced amino acid sets encoded by NDT (C, D, F, G, H, I, L, N, R, S, V, Y) and NRT (C, D, G, H, N, R, S, Y) codons. The activity of the best mutant identified using the respective strategy is displayed relative to the best double mutant among all 400 tested for each ArM reaction. Grey bars represent the mean across all “hits” for the five reactions as identified by the different strategies.

Next, we compared the full-factorial screening approached as pursued herein to other common enzyme engineering strategies. To this end, we used our screening results to analyze which variants would have been identified using such heuristics, and how their activity compares to the most active variant identified in our exhaustive screening. As highlighted in Figure 4c, screening only single mutants (S112X K121 or S112 K121X) would have led to “hits” with activities ranging from 26 to 65 % relative to the most active double mutant for the respective ArM reaction. Subsequent combination of the most beneficial single mutations would have led to an activity increase in two cases, but a decrease in three cases. This suggests that there are interactions between the two amino acid positions 112 and 121 that lead to non-additive effects on the overall activity^46^. Indeed, further analysis revealed that between 24 and 37 % of the observed variance can be attributed to interactions between the two positions (Supplementary Table 1). Notably, the activity of the best mutants identified in our screening can only be explained by considering non-additive effects (Supplementary Fig. 9).

Iterative saturation mutagenesis (ISM) relies on the sequential randomization of positions, while using the best variant from the previous round as a starting point for the next^47^. In this way, it can leverage non-additive interactions to some extent while keeping the experimental effort limited. Indeed, this strategy would have been more successful than the simple combination of single mutations and would have led to the discovery of mutants at or near the local activity optimum for two out of five reactions. Nonetheless, this demonstrates that a full-factorial library enables the discovery of mutants that would likely be missed using other strategies as they are the result of non-trivial interactions. In addition, note that while sequential strategies require the analysis of fewer variants, they do not provide universally applicable libraries, which would be a substantial disadvantage for a general ArM screening platform as proposed here.

Unfortunately, full-factorial screening quickly becomes intractable if more positions are to be diversified. An established strategy to decrease library size is to use reduced amino acid sets, such as the twelve amino acids encoded by NDT codons, which still cover all major classes of amino acid chemistries and lack redundant or stop codons^48^. Our analysis shows that this strategy would have been more successful than those discussed above, reaching consistently high activities for all ArM reactions. This observation is in line with the fact that these twelve amino acids cover the clusters identified in our previous analysis well (Fig. 4b). For this reason, it seems promising to screen combinatorial libraries of Sav mutants based on such a reduced amino acid set. Further reductions of the amino acid set are possible (for example eight amino acids encoded by NRT), but at the cost of a reduced probability of finding highly active mutants.

## Discussion

The results presented in this study highlight that Sav-based ArMs can be tailored to catalyze various new-to-nature reactions. The activity landscapes of ArMs are distinct for each reaction, underscoring the need for flexible enzyme engineering strategies that can be easily adapted to new transformations. The Sav 112X 121X library along with the automation-compatible, periplasmic screening workflow represent the first platform for systematic ArM engineering to this end. Using this approach, we readily identified improved ArMs for five biorthogonal reactions. Importantly, the activity enhancements compare favorably to previous efforts of optimizing Sav-based ArMs^34,35^. Similarly, the “hits” appear to be more active than variants we identified previously for the same reactions. For instance, several studies had previously described allylic deallylases that uncage allylcarbamate-protected substrates. The most extensive engineering study of these was a screening of 80 surface-displayed variants with mutations at positions 112 and 121, yielding the double mutants 112M 121A and 112Y 121S as the most active variants for the uncaging of allylcarbamate-protected amino coumarin **3** (**II**)^35^. While both of these mutants proofed to be highly active also in our periplasmic screen, several others displayed even higher activities, reaching an improvement over wild type of up to fifteen-fold, compared to nine- and ten-fold for the previously identified variants.

The recorded data on ArM activity and cellular Sav levels indicate that differences in the expression level of Sav mutants do not affect the screening performance of our platform, as an excess of biotin-binding sites is consistently ensured. This notion is corroborated by the observation that the best ArM mutants displayed an increased specific activity *in vitro*. At the same time, the large excess of binding sites points to a substantial potential to increase the cell-specific ArM activity for applications in biocatalysis, which could be achieved by increasing the concentration of cofactor and/or improving its uptake.

All reactions tested in the context of this study were found to be compatible with whole-cell biotransformations under mild reaction conditions, which is an important prerequisite for advanced applications in the context of synthetic biology^5^. In this regard, the indole-producing ArMs (reactions **III** and **IV**) are of particular interest, as a variety of applications can be imagined based on this metabolic intermediate and signaling molecule.

The combinatorial library focused on two crucial residues proofed to be a powerful tool for the engineering of ArMs, particularly when pursuing multiple catalytic activities in parallel from a common starting point. The combination of such libraries with the screening workflow presented herein enables the rapid discovery of active ArMs for any new reaction, potentially as a universal initial step followed by more extensive engineering campaigns. For the latter, the results from this study provide valuable implications such as the indicated high potential of reduced amino acid sets in the context of ArMs. Combined with lab automation, such amino acid sets render larger screening campaigns involving more amino acid positions feasible. For much larger numbers of targeted positions in Sav, ISM, potentially in combination with reduced amino acid alphabets, appears to be a promising strategy as it offers a good trade-off between experimental effort and the probability of finding highly active mutants. Lastly, as an alternative to the exhaustive search pursued here, the screening platform may be used to perform undersampling of libraries with a much larger theoretical diversity. In combination with powerful machine learning techniques, which have been ascribed a high potential for enzyme engineering in general^49^, this would enable smartly guided and efficient directed evolution approaches for ArMs.

## Acknowledgments

This work was kindly supported by the NCCR “Molecular Systems Engineering” and the European Commission (project: Madonna; grant no. 766975). TRW thanks the SNF (grant no. 200020_182046) for generous support. SP and TV acknowledge support by the SNF (grant no. 31003A_179521). The authors kindly thank Valerio Sabatino, Fabian Schwizer, Yoann Cotelle and Jaicy Vallapurackal for their help with substrate and cofactor synthesis, and Gregor Schmidt for technical support regarding lab automation.

## Author Contributions

MJ and TRW conceived and supervised the study. SP advised the project. TV carried out biological and screening experiments and analyzed the data. FC and MP developed the gold-catalyzed reactions *in vitro* and synthesized the corresponding substrates and cofactors. TV, MJ, SP and TRW discussed the data. TV and MJ wrote the manuscript with input from all authors.

## Methods

### Chemicals and reagents

Unless stated otherwise, chemicals were obtained from Sigma-Aldrich, Acros or Fluorochem. Primers were synthesized by Sigma-Aldrich and enzymes for molecular cloning were obtained from New England Biolabs.

### Cloning of Sav library

The Sav library was created based on a previously described expression plasmid that contains a T7-tagged Sav gene with an N-terminal *ompA* signal peptide for export to the periplasm under control of the T7 promoter in a pET30b vector (Addgene #138589)^34^. Positions 112 and 121 were randomized using the codon set described by Tang *et al*.^50^ The plasmid was amplified in two fragments using either primer 1 and a mix of primers 2-5 (molar ratio of 12:6:1:1) or primer 10 and a mix of primers 6-9 (molar ratio of 12:6:1:1, see Supplementary Tab. 2 for primer sequences). PCRs were carried out using Q5 polymerase. Template plasmid was digested using DpnI and the PCR products were purified using a PCR purification kit (Sigma-Aldrich). Subsequently, the fragments were joined by Gibson assembly and used to transform chemically competent BL21-Gold(DE3) cells (Agilent Technologies). Individual clones were sequence-verified by Sanger sequencing (Microsynth AG). Having identified 250 out of 400 possible distinct double mutants this way, a second pool containing only the 150 missing variants was cloned. To this end, sequence-verified plasmids from the first library generation step were used as PCR templates and 40 fragments, each containing a single amino acid exchange at position 112 or 121, were obtained by amplification with primers 1 and 12 or 10 and 11 for positions 112 and 121, respectively. Following DpnI digest and purification, these fragments were added to Gibson assembly reactions in combinations suitable for obtaining pools of missing variants. More specifically, this was achieved by adding one fragment with a desired mutation at position 112 per reaction, along with multiple fragments for position 121. Again, individual clones were sequenced after transformation. Eventually, 36 remaining variants were cloned individually by assembling the corresponding fragments. Refer to Supplementary Figure 1 for an overview of the library generation.

### Sav expression in 96-well plates

96 deep-well plates were filled with 500 µL LB (+ 50 mg L^-1^ kanamycin) per well. Cultures were inoculated from glycerol stocks and grown overnight at 37 °C and 300 rpm in a Kuhner LT-X shaker (50 mm shaking diameter). 20 µL per culture were used to inoculate expression cultures in 1 mL LB with kanamycin. These cultures were grown at 37 °C and 300 rpm for 1.5 h. At this point, the plates were placed at room temperature for 20 min and subsequently Sav expression was induced by addition of 50 µM isopropyl β-D-1-thiogalactopyranoside (IPTG). Expression was carried out at 20 °C and 300 rpm for an additional 20 h.

### Quantification of biotin-binding sites

For measurement of Sav expression levels, the OD_600_ of the cultures was determined in a plate reader (Infinite M1000 PRO, Tecan Group AG) using 50 µL samples diluted with an equal volume of phosphate-buffered saline (PBS). The remaining cultures were then centrifuged (3,220 rcf, 20 °C, 10 min) and the pellets were resuspended in 250 µL lysis buffer (50 mM Tris, 150 mM NaCl, 5 mM EDTA, 1 g L^-1^ lysozyme, pH 7.4). After 30 min incubation at room temperature, the cell suspensions were subjected to three freeze-thaw cycles. Afterwards, 150 µL of DNaseI buffer (50 mM Tris, 150 mM NaCl, 2 units mL^-1^ DNaseI, pH 7.4) were added and plates were incubated at 37 °C for 45 min before centrifugation (4800 rcf, 20 °C, 20 min). Subsequently, the concentration of biotin-binding sites in the supernatant was determined using a modified version of the assay described by Kada *et al*.^40^, which relies on the quenching of the fluorescence of a biotinylated fluorophore upon binding to Sav. Specifically, 190 µL binding site buffer (1 µM biotin-4-fluorescein, 0.1 g L^-1^ bovine serum albumin in PBS) were mixed with 10 µL supernatant or purified Sav standard. After incubation at room temperature for 90 min, the fluorescence intensity was measured (excitation at 485 nm, emission at 525 nm) and a calibration curve produced with purified Sav was used to calculate the concentration of biotin-binding sites in each sample.

### Synthesis of cofactors and substrates

Substrates **1**^34^, **3**^35^, **5**^51^ and cofactors **(Biot-NHC)Ru**^52^ and **(Biot-HQ)CpRu**^35^ were prepared according to reported procedures. Substrate **7** was obtained from Sigma-Aldrich. A detailed description of the synthesis of **(Biot-NHC)Au1, (Biot-NHC’)Au2**, substrate **8** and product **9** is available in the Supplementary Notes.

### Whole-cell screening

Following the expression of Sav mutants in deep-well plates, the OD_600_ of the cultures was determined in a plate reader using 50 µL samples diluted with an equal volume of PBS. Afterwards, the plates were centrifuged (3,220 rcf, 15 °C, 10 min), the supernatant was discarded and the pellets were resuspended in 400 µL incubation buffer containing the respective reaction-specific cofactor. The composition of the incubation buffer varied for metathesis (2 µM **(Biot-NHC)Ru** in 50 mM Tris, 0.9 % (w/v) NaCl, pH 7.4), deallylation (5 µM **(Biot-HQ)CpRu** in 50 mM MES, 0.9 % NaCl, pH 6.1), hydroamination (10 µM **(Biot-NHC)Au1** in 50 mM MES, 0.9 % NaCl, 5 mM diamide, pH 6.1) and hydroarylation (10 µM **(Biot-NHC’)Au2** in 50 mM MES, 0.9 % NaCl, 5 mM diamide, pH 5) reactions. The compositions of different buffers are summarized in Supplementary Table 3. Cells were incubated with the cofactor for 1 h at 15 °C and 300 rpm. Afterwards, plates were centrifuged (2,000 rcf, 15 °C, 10 min), the supernatant was discarded and the pellets were resuspended in 500 µL of the respective incubation buffer lacking cofactor to remove unbound cofactor. Following another centrifugation step, cell pellets were resuspended in 200 µL reaction buffer containing the respective substrate. The composition of this reaction buffer varied for metathesis (5 mM **1** in 100 mM sodium acetate, 0.5 M MgCl_2_, pH 4), deallylation (500 µM **3** or **5** in 50 mM MES, 0.9 % NaCl, pH 6.1), hydroamination (5 mM **7** in 50 mM MES, 0.9 % NaCl, 5 mM diamide, pH 6.1) and hydroarylation (5 mM **9** in 50 mM MES, 0.9 % NaCl, 5 mM diamide, pH 5) reactions (Supplementary Tab. 3). Reactions were performed at 37 °C and 300 rpm for 20 h before determining the product concentration. Each 96-well plate contained 80 mutants to be tested along with four replicates of cells expressing wild-type Sav, Sav SL, Sav MA and cells lacking Sav as controls. To account for differences in cell density and plate-to-plate variations, the product concentrations were divided by the OD_600_ of the culture and normalized to the mean of the cell-specific product concentrations measured for the Sav SL (metathesis, hydroamination and hydroarylation) or Sav MA (deallylation) controls on the respective plate. These mutants had been identified as active in preliminary experiments. All variants were tested at least in biological duplicates.

### Lab automation

The steps required for incubation of cells with cofactor, washing, substrate addition and OD_600_ measurement were implemented on two connected Tecan Freedom EVO 200 Robots, equipped with a Kuhner deep-well plate shaker for incubations, a Rotanta 46 RSC centrifuge and a Tecan Infinite M200 PRO plate reader for OD_600_ measurement. The automated protocol was used for the deallylation with allyl carbamate coumarin (**II**) and the gold-catalyzed hydroarylation (**V**), while the screenings for the other reactions were performed manually.

### Product quantification by UPLC-MS

To quantify the metathesis product **2**, an extraction was performed by adding 775 µL methanol and 25 µL internal standard (100 µM in methanol) to each 200 µL sample. The samples were incubated for one hour at 20 °C and 300 rpm, followed by centrifugation at 3,220 rcf and 20 °C for 10 min. Subsequently, 50 µL of the supernatant were mixed with 200 µL water and analyzed by UPLC-MS. UPLC analysis was performed using a Waters H-Class Bio using a BEH C18 1.7 μM column and a flow rate of 0.6 ml min^−1^ (eluent A, 0.1 % formic acid in water; eluent B, 0.1% formic acid in acetonitrile; gradient at 0 min: 90 % A, 10 % B; at 0.5 min: 90 % A, 10 % B; at 2.5 min: 10 % A, 90 % B; at 3.5 min: 90 % A, 10 % B; at 4.5 min: 90 % A, 10 % B). Peak integration for SIR (single ion recording) were used for quantification, and concentrations of the metathesis product (retention time of 1.0 min ± 0.25 min) were determined on the basis of a standard curve with the ring-closed product in the presence of a fixed concentration of the nonadeuteraded product (retention time of 1.0 min ± 0.25 min).

### Fluorescence measurements

The fluorescent product **4** was quantified by measuring the fluorescence intensity at an excitation of 394 nm and emission of 460 nm. Product **9** was excited at 390 nm and measured at 488 nm. Measurements were carried out in black 96-well plates in an Infinite M1000 PRO plate reader.

### Kovac’s assay

Indole was quantified using the photometric Kovac’s assay (adapted from Piñero-Fernandez *et al*.^53^). For measurements in culture supernatant, plates were centrifuged (3,220 rcf, 20 °C, 10 min) and 110 µL supernatant was mixed with 165 µL Kovac’s reagent (50 g L^-1^ 4- (dimethylamino)benzaldehyde, 710 g L^-1^ isoamylic alcohol, 240 g L^-1^ hydrochloric acid) in a separate plate. After 5 min incubation, these plates were centrifuged (3,220 rcf, 20 °C, 10 min). Subsequently, 75 µL of the upper phase were transferred to a new transparent plate and the absorbance at 540 nm was measured in a plate reader (Infinite M1000 PRO). The same procedure (omitting the first centrifugation step) was applied to measure indole after *in vitro* experiments. A standard curve was then used to determine indole concentrations.

### Sav expression for purification

A single colony of BL21-Gold(DE3) harboring a plasmid from the previously described library for periplasmic expression of the desired Sav variant was used to inoculate a starter culture (4 mL LB with 50 mg L^-1^ kanamycin), which was grown overnight at 37 °C and 200 rpm. On the following day, 100 mL LB with kanamycin in a 500 mL flask were inoculated to an OD_600_ of 0.01. The culture was grown at 37 °C and 200 rpm until it reached an OD_600_ of 0.5. At this point, the flask was placed at room temperature for 20 min and 50 µM IPTG were added to induce Sav expression. Expression was performed at 20 °C and 200 rpm overnight, and cells were harvested by centrifugation (3,220 rcf, 4 °C, 15 min). Pellets were stored at -20 °C until purification.

### Sav purification

Cell pellets were resuspended in lysis buffer (50 mM Tris, 150 mM NaCl, 1 g L^-1^ lysozyme, pH 7.4) so as to reach a final OD_600_ of 10. After 30 min incubation at room temperature, cell suspensions were subjected to three freeze-thaw cycles. Subsequently, nucleic acids were digested by addition of 1 µL benzonase (Merck KGaA) and 10 µL 1 M MgSO_4_ followed by incubation at room temperature for 10 min. After centrifugation, the supernatant was transferred to a new tube and mixed with carbonate buffer (50 mM ammonium bicarbonate, 500 mM NaCl, pH 11) in a ratio of 2:3. 500 µL iminobiotin beads were added and the tubes were incubated at room temperature with shaking (120 rpm) for 1 h. Afterwards, the beads were washed twice with 15 mL carbonate buffer and resuspended in 2 mL acetate buffer (50 mM ammonium acetate, 500 mM NaCl, pH 4). After 20 min of incubation at room temperature with shaking, the tubes were centrifuged and the supernatant was dialyzed (SnakeSkin dialysis tubing with 3.5 kDa molecular weight cut-off, Thermo Fisher Scientific) against 1 L of the desired buffer (Supplementary Table 4) for 24 h at room temperature with one buffer exchange.

### *In vitro* catalysis

*In vitro* reactions were performed with 2.5 µM purified Sav (tetrameric; corresponding to 10 µM biotin-binding sites) in a volume of 200 µL in glass vials. Cofactor and substrate concentrations as well as buffer conditions varied between reactions and are listed in Supplementary Table 2. Reactions were carried out at 37 °C and 200 rpm for 20 h.

### Data analysis

Data were analyzed using R 4.0.2^54^. For clustering and calculating amino acid effects, activity values were standardized by subtracting the mean and dividing by the standard deviation of all activity values for the respective reaction. Hierarchical clustering was done by calculating the Euclidian distances between all standardized activity values of variants harboring the respective amino acid at position 112 or 121 (20 values per position and reaction, amounting to 200 values in total) and clustering these using the hclust function^54^ with complete linkage. The activity of all 400 mutants was further analyzed by two-way analysis of variance (ANOVA). To this end, position 112 and 121 were treated as explanatory variables (with 20 levels corresponding to the canonical amino acids). For analyzing the variance explained by individual factors, an interaction term was included, whereas it was omitted in order to generate a purely additive model.

### Data and code availability

All data and code required to reproduce the figures and analyses presented in the main text will be made publicly accessible upon publication.

## References

1. Lewis, J. C. Artificial metalloenzymes and metallopeptide catalysts for organic synthesis. ACS Catal. 3, 2954–2975 (2013).

2. Rosati, F. & Roelfes, G. Artificial metalloenzymes. ChemCatChem 2, 916–927 (2010).

3. Bornscheuer, U. T. The fourth wave of biocatalysis is approaching. Philos. Trans. R. Soc. A Math. Phys. Eng. Sci. 376, 20170063 (2018).

4. Hammer, S. C., Knight, A. M. & Arnold, F. H. Design and evolution of enzymes for non-natural chemistry. Curr. Opin. Green Sustain. Chem. 7, 23–30 (2017).

5. Vornholt, T. & Jeschek, M. The quest for xenobiotic enzymes - from new enzymes for chemistry to a novel chemistry of life. ChemBioChem (2020). doi: 10.1002/cbic.202000121

6. Kan, S. B. J., Lewis, R. D., Chen, K. & Arnold, F. H. Directed evolution of cytochrome c for carbon–silicon bond formation: Bringing silicon to life. Science 354, 1048–1051 (2016).

7. Tinoco, A., Steck, V., Tyagi, V. & Fasan, R. Highly diastereo- and enantioselective synthesis of trifluoromethyl-substituted cyclopropanes via myoglobin-catalyzed transfer of trifluoromethylcarbene. J. Am. Chem. Soc. 139, 5293–5296 (2017).

8. Song, W. J. & Tezcan, F. A. A designed supramolecular protein assembly with in vivo enzymatic activity. Science 346, 1525–1528 (2014).

9. Studer, S. et al. Evolution of a highly active and enantiospecific metalloenzyme from short peptides. Science 362, 1285–1288 (2018).

10. Drienovská, I. et al. Design of an enantioselective artificial metallo-hydratase enzyme containing an unnatural metal-binding amino acid. Chem. Sci. 8, 7228–7235 (2017).

11. Cangelosi, V. M., Deb, A., Penner-Hahn, J. E. & Pecoraro, V. L. A de novo designed metalloenzyme for the hydration of CO2. Angew. Chem. Int. Ed. 53, 7900–7903 (2014).

12. Key, H. M., Dydio, P., Clark, D. S. & Hartwig, J. F. Abiological catalysis by artificial haem proteins containing noble metals in place of iron. Nature 534, 534–537 (2016).

13. Yang, H. et al. Evolving artificial metalloenzymes via random mutagenesis. Nat. Chem. 10, 318–324 (2018).

14. Mirts, E. N., Petrik, I. D., Hosseinzadeh, P., Nilges, M. J. & Lu, Y. A designed heme-[4Fe-4S] metalloenzyme catalyzes sulfite reduction like the native enzyme. Science 361, 1098–1101 (2018).

15. Raines, D. J. et al. Redox-switchable siderophore anchor enables reversible artificial metalloenzyme assembly. Nat. Catal. 1, 680–688 (2018).

16. Eda, S. et al. Biocompatibility and therapeutic potential of glycosylated albumin artificial metalloenzymes. Nat. Catal. 2, (2019).

17. Dydio, P. et al. An artificial metalloenzyme with the kinetics of native enzymes. Science 354, 102–106 (2016).

18. Oohora, K., Kihira, Y., Mizohata, E., Inoue, T. & Hayashi, T. C(sp3)-H bond hydroxylation catalyzed by myoglobin reconstituted with manganese porphycene. J. Am. Chem. Soc. 135, 17282–17285 (2013).

19. Chevalley, A., Cherrier, M. V., Fontecilla-Camps, J. C., Ghasemi, M. & Salmain, M. Artificial metalloenzymes derived from bovine β-lactoglobulin for the asymmetric transfer hydrogenation of an aryl ketone – synthesis, characterization and catalytic activity. Dalt. Trans. 43, 5482–5489 (2014).

20. Lopez, S. et al. A mechanistic rationale approach revealed the unexpected chemoselectivity of an artificial Ru-dependent oxidase: A dual experimental/theoretical approach. ACS Catal. 10, 5631–5645 (2020).

21. Burke, A. J. et al. Design and evolution of an enzyme with a non-canonical organocatalytic mechanism. Nature 570, 219–223 (2019).

22. Emmanuel, M. A., Greenberg, N. R., Oblinsky, D. G. & Hyster, T. K. Accessing non-natural reactivity by irradiating nicotinamide-dependent enzymes with light. Nature 540, 414–417 (2016).

23. Biegasiewicz, K. F. et al. Photoexcitation of flavoenzymes enables a stereoselective radical cyclization. Science 364, 1166–1169 (2019).

24. Heinisch, T. & Ward, T. R. Artificial metalloenzymes based on the biotin-streptavidin technology: challenges and opportunities. Acc. Chem. Res. 49, 1711–1721 (2016).

25. Nödling, A. R. et al. Reactivity and selectivity of iminium organocatalysis improved by a protein host. Angew. Chem. Int. Ed. 57, 12478–12482 (2018).

26. Hassan, I. S. et al. Asymmetric δ-lactam synthesis with a monomeric streptavidin artificial metalloenzyme. J. Am. Chem. Soc. 141, 4815–4819 (2019).

27. Zeymer, C. & Hilvert, D. Directed evolution of protein catalysts. Annu. Rev. Biochem. 87, 131–157 (2018).

28. Zhang, R. K., Romney, D. K., Kan, S. B. J. & Arnold, F. H. Directed evolution of artificial metalloenzymes: bridging synthetic chemistry and biology. in Artificial metalloenzymes and metalloDNAzymes in catalysis 137–170 (Wiley-VCH Verlag GmbH & Co. KGaA, 2018). doi: 10.1002/9783527804085.ch5

29. Reetz, M. T. Directed evolution of artificial metalloenzymes: a universal means to tune the selectivity of transition metal catalysts? Acc. Chem. Res. (2019). doi: 10.1021/acs.accounts.8b00582

30. Markel, U., Sauer, D. F., Schiffels, J., Okuda, J. & Schwaneberg, U. Towards the evolution of artificial metalloenzymes—a protein engineer’s perspective. Angew. Chem. Int. Ed. 58, 4454–4464 (2019).

31. Turner, N. J. Directed evolution of enzymes for applied biocatalysis. Trends Biotechnol. 21, 474–478 (2003).

32. Creus, M. et al. X-ray structure and designed evolution of an artificial transfer hydrogenase. Angew. Chem. Int. Ed. 47, 1400–1404 (2008).

33. Hyster, T. K., Knorr, L., Ward, T. R. & Rovis, T. Biotinylated Rh(III) complexes in engineered streptavidin for accelerated asymmetric C-H activation. Science 338, 500–503 (2012).

34. Jeschek, M. et al. Directed evolution of artificial metalloenzymes for in vivo metathesis. Nature 537, 661–665 (2016).

35. Heinisch, T. et al. E. coli surface display of streptavidin for directed evolution of an allylic deallylase. Chem. Sci. 9, 5383–5388 (2018).

36. Grimm, A. R. et al. A whole cell E. coli display platform for artificial metalloenzymes: poly(phenylacetylene) production with a rhodium-nitrobindin metalloprotein. ACS Catal. 8, 2611–2614 (2018).

37. Donnelly, A. E., Murphy, G. S., Digianantonio, K. M. & Hecht, M. H. A de novo enzyme catalyzes a life-sustaining reaction in Escherichia coli. Nat. Chem. Biol. 14, 253–255 (2018).

38. Wilson, Y. M., Dürrenberger, M., Nogueira, E. S. & Ward, T. R. Neutralizing the detrimental effect of glutathione on precious metal catalysts. J. Am. Chem. Soc. 136, 8928–8932 (2014).

39. Jeschek, M., Panke, S. & Ward, T. R. Periplasmic screening for artificial metalloenzymes. in Methods in Enzymology 580, 539–556 (Elsevier Inc., 2016).

40. Kada, G., Kaiser, K., Falk, H. & Gruber, H. J. Rapid estimation of avidin and streptavidin by fluorescence quenching or fluorescence polarization. Biochim. Biophys. Acta 1427, 44–8 (1999).

41. Tokuriki, N. & Tawfik, D. S. Stability effects of mutations and protein evolvability. Curr. Opin. Struct. Biol. 19, 596–604 (2009).

42. Völker, T., Dempwolff, F., Graumann, P. L. & Meggers, E. Progress towards bioorthogonal catalysis with organometallic compounds. Angew. Chem. Int. Ed. 53, 10536–10540 (2014).

43. Völker, T. & Meggers, E. Chemical activation in blood serum and human cell culture: improved ruthenium complex for catalytic uncaging of alloc-protected amines. ChemBioChem 18, 1083–1086 (2017).

44. Do, J. H., Kim, H. N., Yoon, J., Kim, J. S. & Kim, H. J. A rationally designed fluorescence turn-on probe for the gold(III) ion. Org. Lett. 12, 932–934 (2010).

45. Sasmal, P. K., Streu, C. N. & Meggers, E. Metal complex catalysis in living biological systems. Chem. Commun. 49, 1581–1587 (2013).

46. Reetz, M. T. The importance of additive and non-additive mutational effects in protein engineering. Angew. Chem. Int. Ed. 52, 2658–2666 (2013).

47. Reetz, M. T. & Carballeira, J. D. Iterative saturation mutagenesis (ISM) for rapid directed evolution of functional enzymes. Nat. Protoc. 2, 891–903 (2007).

48. Reetz, M. T., Kahakeaw, D. & Lohmer, R. Addressing the numbers problem in directed evolution. ChemBioChem 9, 1797–1804 (2008).

49. Yang, K. K., Wu, Z. & Arnold, F. H. Machine-learning-guided directed evolution for protein engineering. Nat. Methods 16, 687–694 (2019).

50. Tang, L. et al. Construction of ‘small-intelligent’ focused mutagenesis libraries using well-designed combinatorial degenerate primers. Biotechniques 52, 149–158 (2012).

51. Jacquemard, U. et al. Mild and selective deprotection of carbamates with Bu4NF. Tetrahedron 60, 10039–10047 (2004).

52. Kajetanowicz, A., Chatterjee, A., Reuter, R. & Ward, T. R. Biotinylated metathesis catalysts: synthesis and performance in ring closing metathesis. Catal. Letters 144, 373–379 (2014).

53. Piñero-Fernandez, S., Chimerel, C., Keyser, U. F. & Summers, D. K. Indole transport across Escherichia coli membranes. J. Bacteriol. 193, 1793–1798 (2011).

54. R Core Team. R: A language and environment for statistical computing. R Foundation for Statistical Computing (2019). Available at: https://www.r-project.org/.

