## Supplementary Information for "Systematic Engineering of Artificial Metalloenzymes for New-to-Nature Reactions"

#### Supplementary Figures

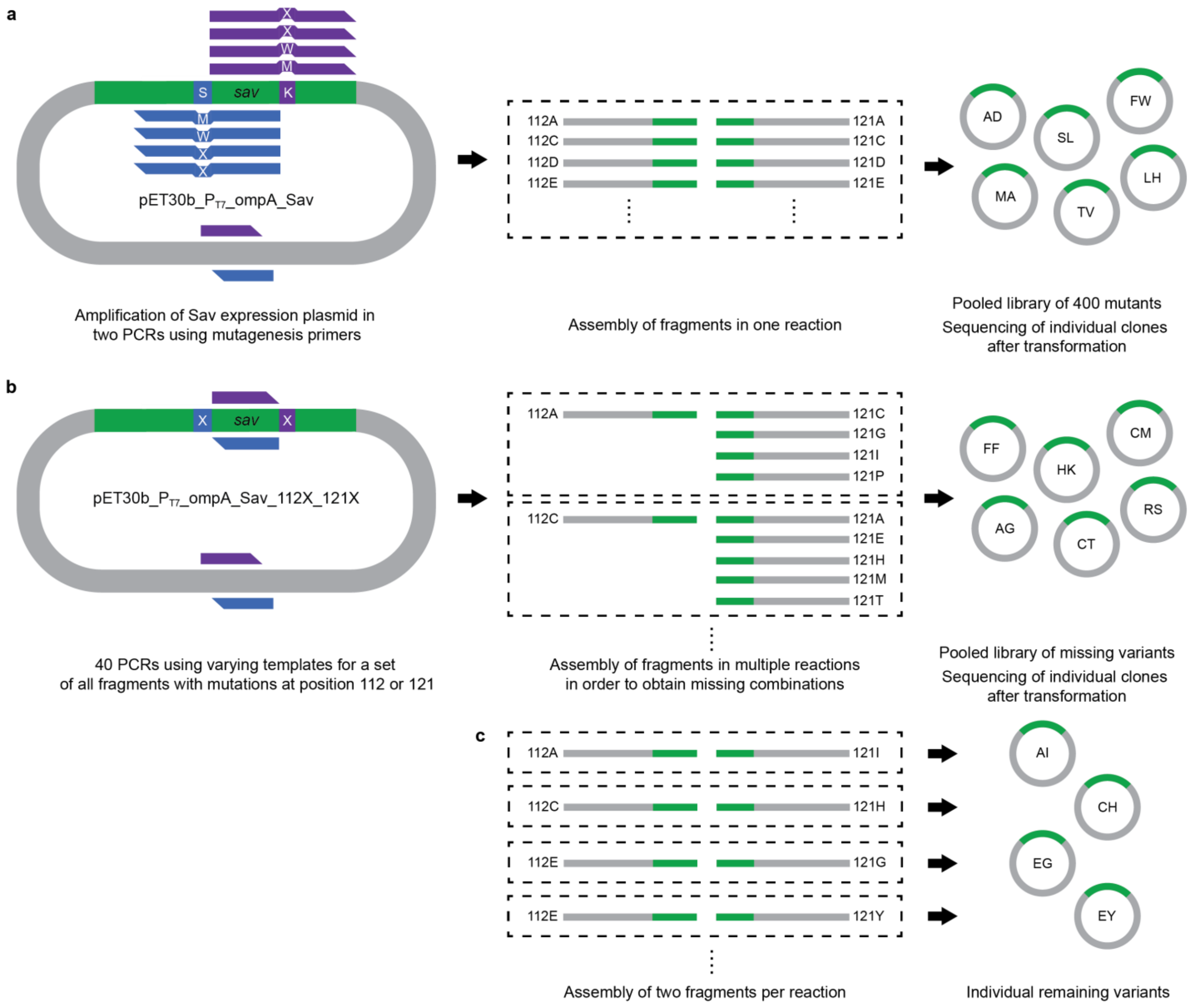


**Supplementary figure 1 | Three-step library-generation strategy. a**, The positions 112 and 121 of Sav in pET30b_PT7_ompA_Sav[34] were randomized using primer mixes to introduce 20 codons encoding all 20 canonical amino acids at the respective positions (mix of four primers). The mixes were obtained by mixing primers containing codons NDT, VMA, TGG and ATG codons in a ratio of 12:6:1:1 as previously described[50]. Two separate PCRs were carried out in order to amplify the whole plasmid in two overlapping fragments (with blue and purple primer sets). The resulting fragments, each containing a mutation at position 112 or 121, were assembled in one Gibson assembly reaction, resulting in a pooled library of 400 Sav variants. Clones from this library were sequenced in order to obtain a defined set of mutants. **b**, After sequencing 672 clones from the 400 mutant library, we cloned a second library containing only variants we had not obtained yet. To this end, 40 PCR products, each containing a single mutation at position 112 or

121, were generated using two primer sets but varying templates (sequence-verified plasmids from the first library). These were assembled in multiple reactions containing one fragment for position 112 and multiple fragments for position 121 in order to generate only missing combinations. As before, individual clones from this library were sequenced. **c**, 36 Sav variants were cloned individually by assembling the previously generated PCR products in separate reactions.

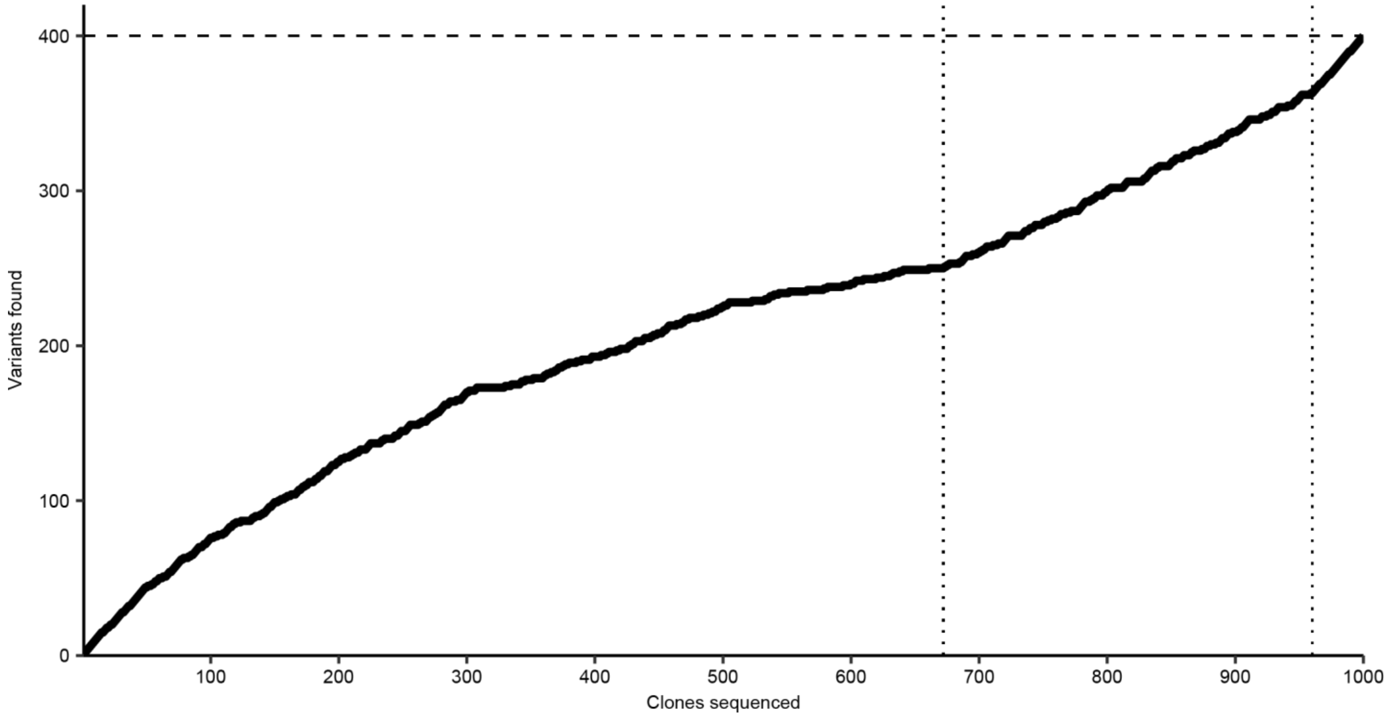


**Supplementary figure 2 | Progress of identifying distinct Sav mutants as a function of the number of sequenced clones.** The vertical lines represent adjustments of the strategy: In the first phase, we sequenced clones from a library created using the "small-intelligent" strategy[50] (Supplementary Fig. 1a). After sequencing 672 clones (leading to the identification of 250 unique variants), we cloned a second library that only contained missing Sav variants (Supplementary Fig. 1b). After sequencing another 288 clones, we cloned the remaining 36 variants individually (Supplementary Fig. 1c) to obtain the full set of 400 Sav 112X 121X mutants.

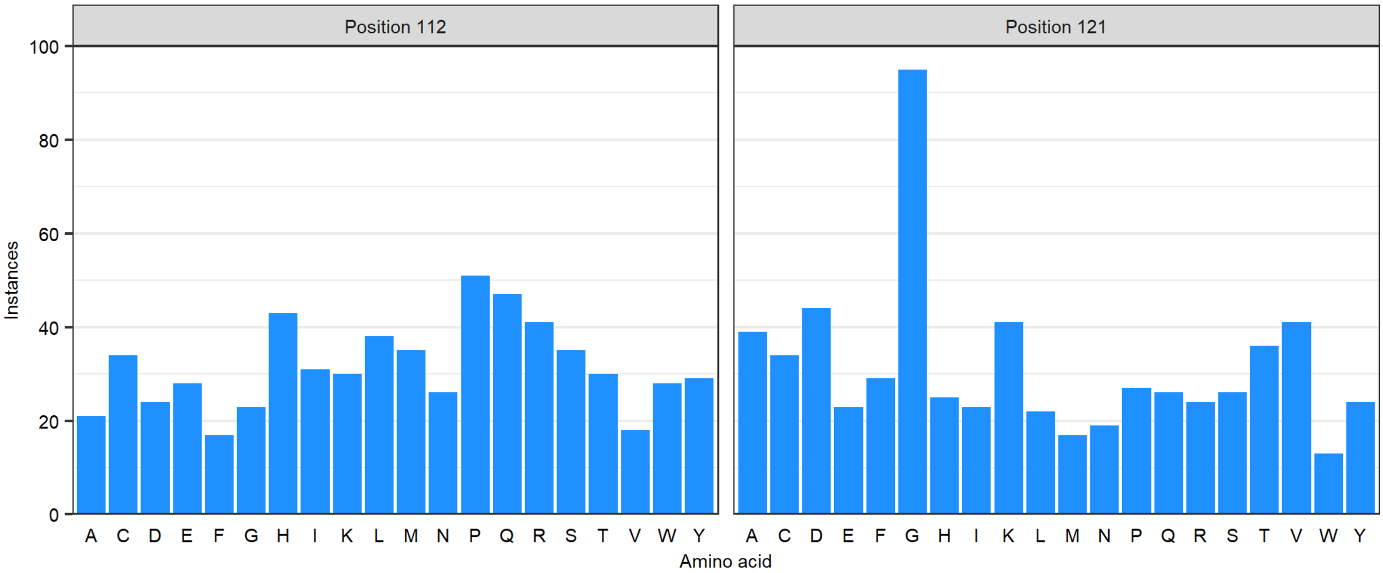


**Supplementary figure 3 | Amino acid distribution in the "small-intelligent" library.** Using the mutagenesis strategy of Tang et al.[50], all 20 amino acids should be present at the same frequency. The sequencing results revealed differences that were typically not larger than two-fold, with the exception of glycine at position 121.

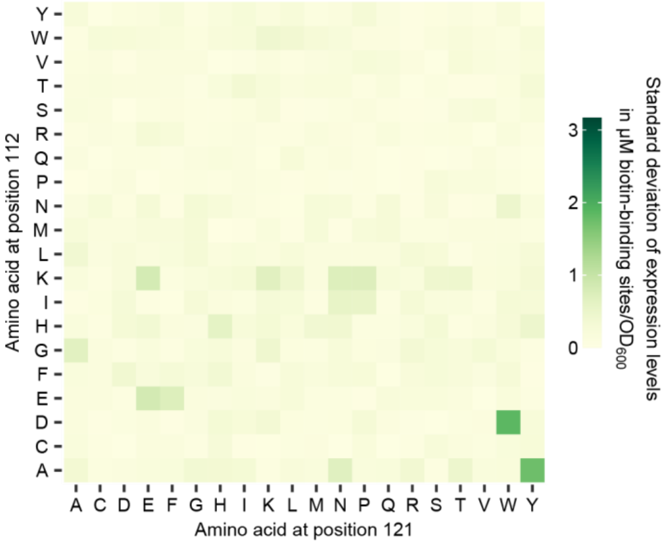


**Supplementary figure 4 | Standard deviation of Sav expression levels.** The concentration of biotin-binding sites was measured in biological triplicate cultures. Displayed values are the corresponding standard deviations and are presented on the same scale as the mean values in Figure 1c.

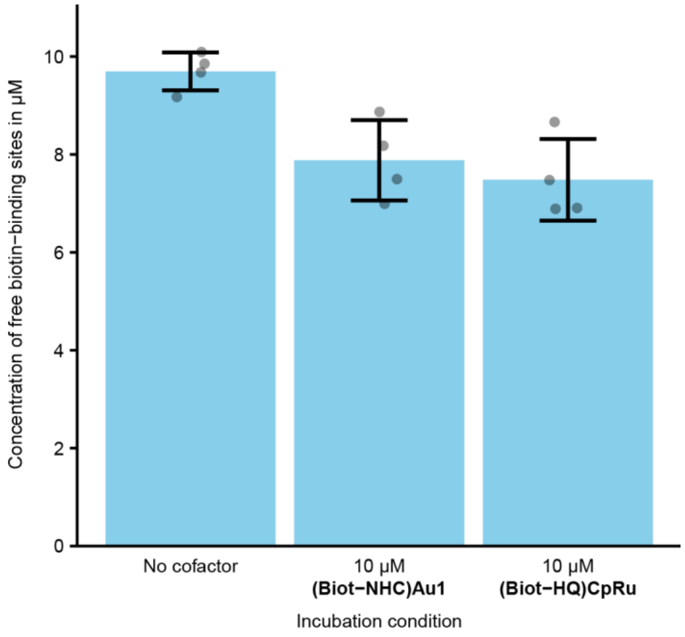


**Supplementary figure 5 | Analysis of cofactor uptake into the periplasm.** Cells expressing periplasmic wild-type Sav were incubated with **(Biot-NHC)Au1**, **(Biot-HQ)CpRu** or without cofactor. Then, the cells were washed and lysed and the concentration of free biotin binding sites was determined[40]. The reduction of free biotin-binding sites for both cofactors relative to the control indicates that below 20 % of the externally added cofactor were taken up.

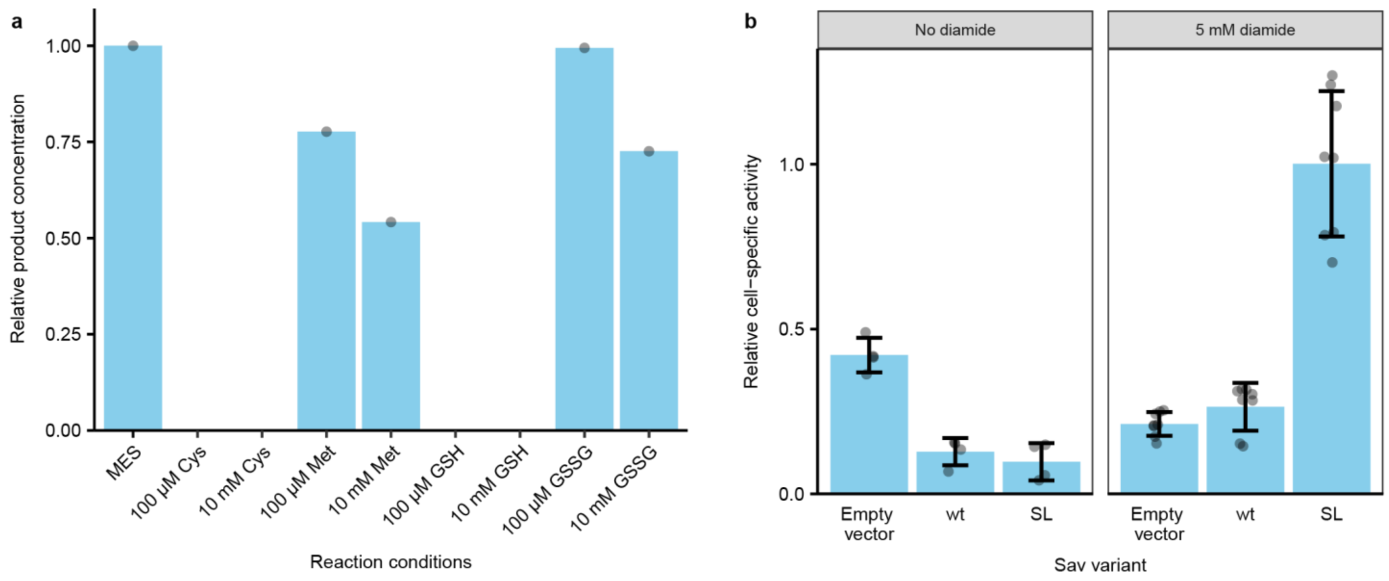


**Supplementary figure 6 | Effect of sulphur-containing metabolites on gold catalysis. a**, *In vitro* reactions with 4 µM Sav biotin binding sites, 2 µM **(Biot-NHC)Au1** and 0.5 mM **7** in MES buffer revealed that thiols such as glutathione (GSH) and cysteine (Cys) have a strong inhibitory effect on the hydroamination reaction. Methionine (Met) and glutathione disulfide (GSSG) had only moderate effects. The y-axis represents the product concentration relative to the control without addition of metabolites (MES). Values are based on single reactions. **b**, Results of periplasmic screening experiments for hydroamination with and without addition of diamide. Active mutants (such as Sav SL) could only be identified reliably when diamide was added. Reactions were performed with four (no diamide) or eight (5 mM diamide) biological replicates. Mean and standard deviation are indicated by bars and error bars, respectively.

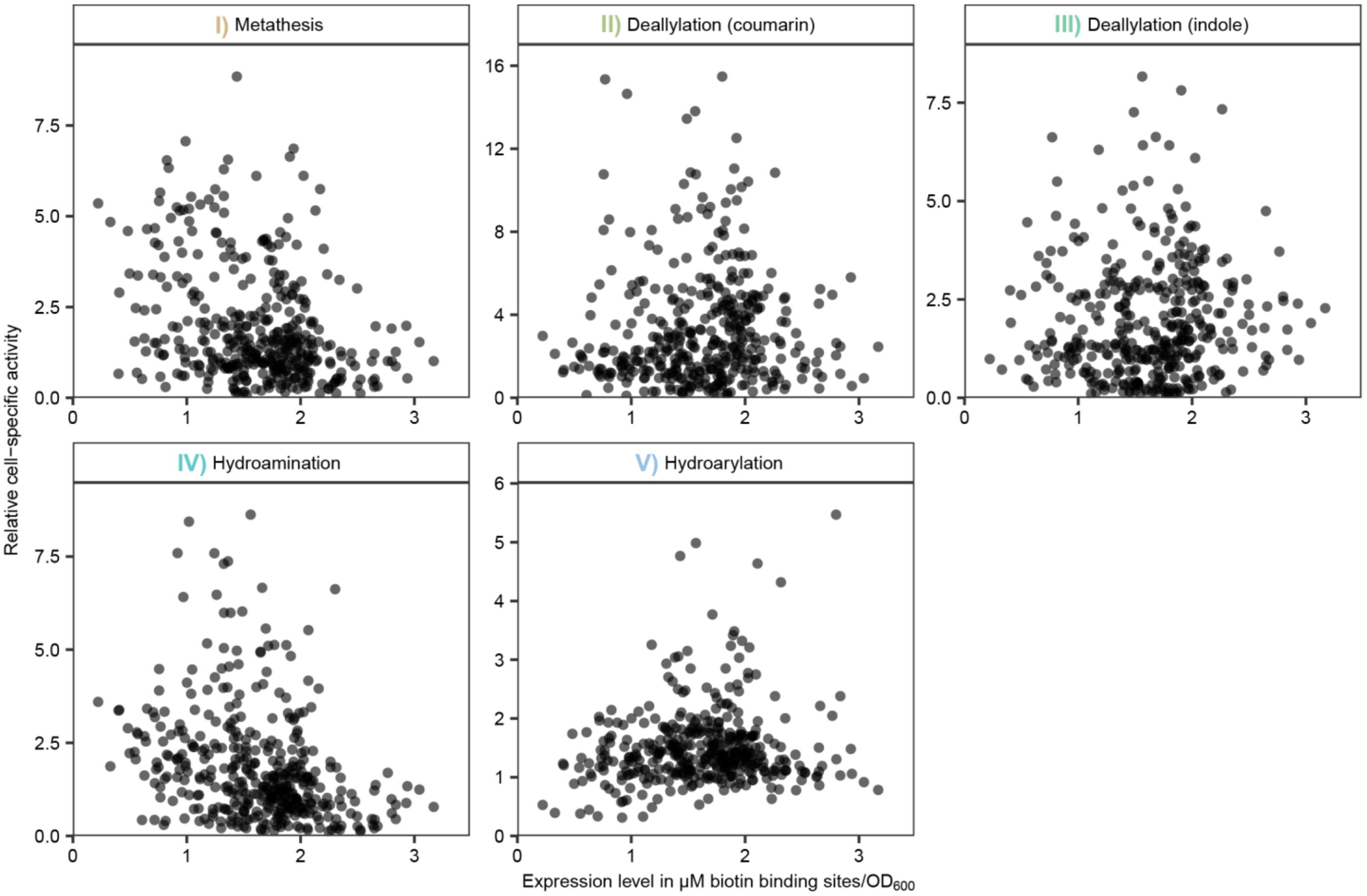


**Supplementary figure 7 | Activity versus expression level.** The cell-specific activity of ArM variants relative to the wild type was plotted against the expression level of Sav (concentration (µM) of biotin-binding sites normalized by the $OD_{600}$). A correlation is not observable for any of the reactions ($R^2 \leq 0.1$).

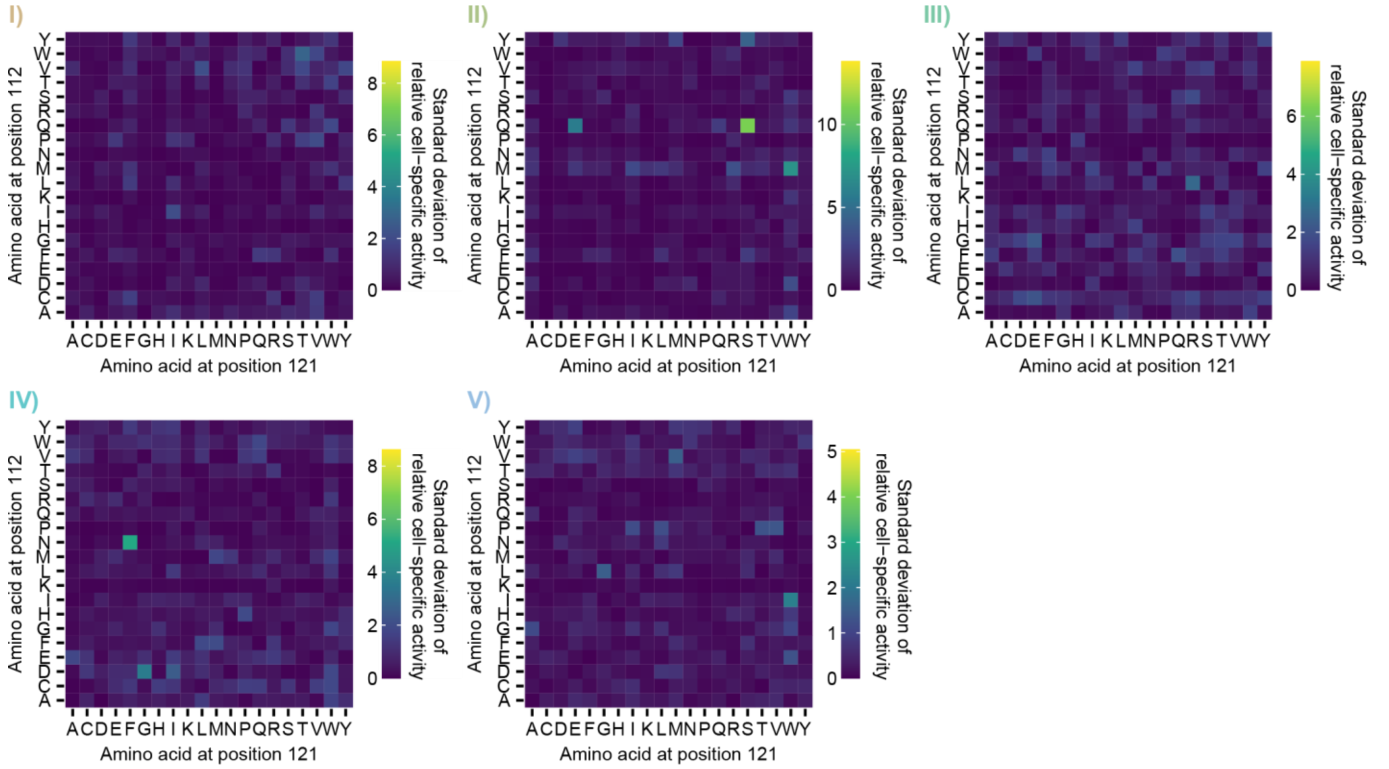


**Supplementary figure 8 | Standard deviation of screening results.** The standard deviation of the relative cell-specific activity of each mutant was determined at least based on biological duplicates. For each reaction, the standard deviation is represented on the same scale as the mean values in Figure 2c**. I)** metathesis, **II)** deallylation (coumarin), **III)** deallylation (indole), **IV)** hydroamination, and **V)** hydroarylation.

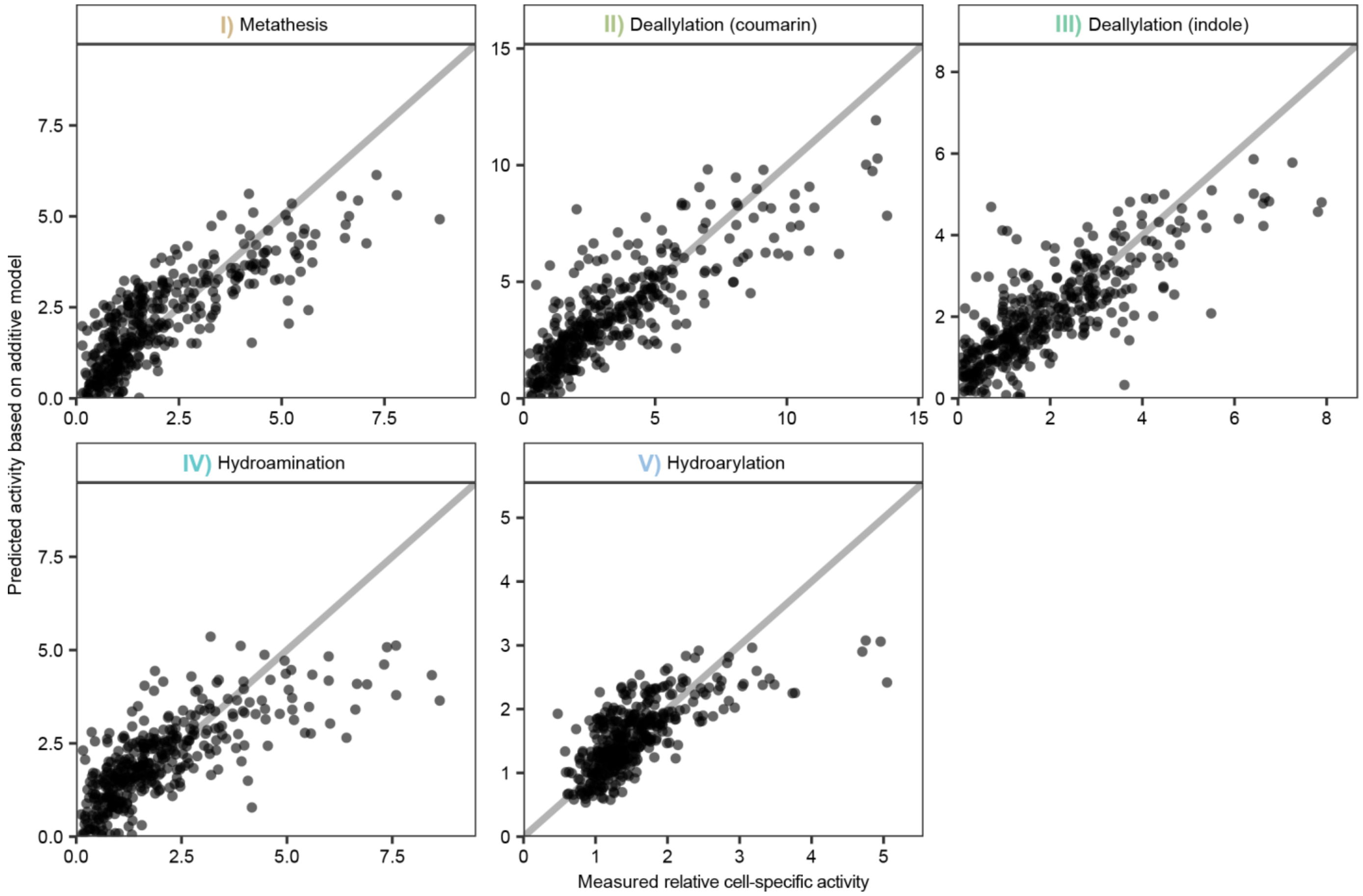


**Supplementary figure 9 | Activity predicted by an additive model versus measured activity.** Analysis of variance (ANOVA) was used to fit a linear model to the measured activities for each reaction. The two Sav positions 112 and 121 were treated as explanatory variables in this model. An interaction term was not included in order to generate a model for the case of purely additive contributions of the amino acids at these positions to the overall activity. The activity values predicted by this model were then plotted against the actual measurements. If the effects of mutations at position 112 and 121 were strictly additive, the points would be located along the diagonal. Points below this line suggest the presence of interactions that increase the activity of the respective mutant, whereas points above the line suggest detrimental interactions. Notably, the most active variants are consistently located below the diagonal, indicating that non-additive interactions are important for obtaining active ArMs.

### Supplementary Tables

**Supplementary table 1 | Relative contribution of position 112, 121 and interactions between the positions to the observed activity.** The screening results for all 400 mutants were analyzed by two-way analysis of variance (ANOVA). Position 112 and position 121 were treated as explanatory variables with 20 levels corresponding to the canonical amino acids. For all reactions, the two positions as well as their interaction factor had a significant influence ($p < 0.001$, F-test). The table summarizes the variance explained (sum of squares of the respective factor divided by total sum of squares) by the individual factors.

| **Reaction** | **Variance explained by position 112 (in %)** | **Variance explained by position 121 (in %)** | **Variance explained by interactions between position 112 and 121 (in %)** | **Residual variance (in %)** |
|---|---|---|---|---|
| Metathesis | 31 | 33 | 27 | 9 |
| Deallylation (coumarin) | 42 | 18 | 24 | 15 |
| Deallylation (indole) | 41 | 14 | 32 | 12 |
| Hydroamination | 24 | 25 | 35 | 16 |
| Hydroarylation | 23 | 19 | 37 | 21 |

**Supplementary table 2 | Sequences of primers used in this study.** Overhangs are displayed in italics and sites for introducing mutations are highlighted in bold.

| # | **Sequence (5' → 3')** |
|---|---|
| 1 | ACAATCTGCTCTGATGCCGCATAG |
| 2 | *CCAGGCGTTGGCCTCGGTGGTGCC**AHN***GGTCAGCAGCCACTGGGTG |
| 3 | *CCAGGCGTTGGCCTCGGTGGTGCC**TKB***GGTCAGCAGCCACTGGGTG |
| 4 | *CCAGGCGTTGGCCTCGGTGGTGCC**CCA***GGTCAGCAGCCACTGGGTG |
| 5 | *CCAGGCGTTGGCCTCGGTGGTGCC**CAT***GGTCAGCAGCCACTGGGTG |
| 6 | *GGCACCACCGAGGCCAACGCCTGG**NDT***TCCACGCTGGTCGGC |
| 7 | *GGCACCACCGAGGCCAACGCCTGG**VMA***TCCACGCTGGTCGGC |
| 8 | *GGCACCACCGAGGCCAACGCCTGG**TGG***TCCACGCTGGTCGGC |
| 9 | *GGCACCACCGAGGCCAACGCCTGG**ATG***TCCACGCTGGTCGGC |
| 10 | GGCTTAACTATGCGGCATCAGAGCAG |
| 11 | GGCACCACCGAGGCCAAC |
| 12 | CCAGGCGTTGGCCTCG |

**Supplementary table 3 | Overview of screening conditions.** Washing buffer was identical to incubation buffer but did not contain cofactor.

| **Reaction** | **Incubation buffer** | **Reaction buffer** |
|---|---|---|
| Metathesis | 2 µM **(Biot-NHC)Ru**<br>50 mM Tris<br>0.9 % (w/v) NaCl<br>pH 7.4 | 5 mM **1**<br>100 mM sodium acetate<br>0.5 M $MgCl_2$<br>pH 4 |
| Deallylation (coumarin) | 5 µM **(Biot-HQ)CpRu**<br>50 mM MES<br>0.9 % (w/v) NaCl<br>pH 6.1 | 500 µM **3**<br>50 mM MES<br>0.9 % (w/v) NaCl<br>pH 6.1 |
| Deallylation (indole) | 5 µM **(Biot-HQ)CpRu**<br>50 mM MES<br>0.9 % (w/v) NaCl<br>pH 6.1 | 500 µM **5**<br>50 mM MES<br>0.9 % (w/v) NaCl<br>pH 6.1 |
| Hydroamination | 10 µM **(Biot-NHC)Au1**<br>50 mM MES<br>0.9 % (w/v) NaCl<br>5 mM diamide<br>pH 6.1 | 5 mM **7**<br>50 mM MES<br>0.9 % (w/v) NaCl<br>5 mM diamide<br>pH 6.1 |
| Hydroarylation | 10 µM **(Biot-NHC')Au2**<br>50 mM MES<br>0.9 % (w/v) NaCl<br>5 mM diamide<br>pH 5 | 5 mM **9**<br>50 mM MES<br>0.9 % (w/v) NaCl<br>5 mM diamide<br>pH 5 |

**Supplementary table 4 | Overview of conditions for *in vitro* catalysis.**

| Reaction | Conditions |
|---|---|
| Metathesis | 10 µM Sav biotin-binding sites<br>0.2 µM **(Biot-NHC)Ru**<br>5 mM **1**<br>100 mM sodium acetate<br>0.5 M $MgCl_2$<br>pH 4 |
| Deallylation (coumarin) | 10 µM Sav biotin-binding sites<br>0.5 µM **(Biot-HQ)CpRu**<br>500 µM **3**<br>50 mM MES<br>0.9 % (w/v) NaCl<br>pH 6.1 |
| Deallylation (indole) | 10 µM Sav biotin-binding sites<br>0.5 µM **(Biot-HQ)CpRu**<br>500 µM **5**<br>50 mM MES<br>0.9 % (w/v) NaCl<br>pH 6.1 |
| Hydroamination | 10 µM Sav biotin-binding sites<br>1 µM **(Biot-NHC)Au1**<br>5 mM **7**<br>50 mM MES<br>0.9 % (w/v) NaCl<br>pH 6.1 |
| Hydroarylation | 10 µM Sav biotin-binding sites<br>1 µM **(Biot-NHC')Au2**<br>5 mM **9**<br>50 mM MES<br>0.9 % (w/v) NaCl<br>10 % (v/v) DMSO<br>pH 5 |

### Supplementary Notes

#### Synthesis of gold cofactors

##### General notes

Procedures for the synthesis of the gold complexes were carried out in dried glassware under a dry nitrogen atmosphere with rigorous exclusion of moisture from reagents and glassware using standard Schlenk techniques or a MBraun Labstar glove box workstation. Dry solvents were directly purchased from Sigma Aldrich or Acros Organics and used without further purification. Water used in the catalytic reactions was purified by Milli-Q Advantage system.

Chemicals were purchased from Sigma Aldrich, Acros Organics, Alfa Aesar or Fluorochem and used without further purification. All catalytic reactions were carried out with non-degassed solvents under air. Temperature was maintained using Thermowatch-controlled heating blocks. Analytical thin-layer chromatography (TLC) was performed on pre-coated Merck silica gel 60 $F_{254}$ plates (0.25 mm) and visualized using UV light. Automated flash column chromatography was carried out on a Biotage Isolera using Silicycle SiliaFlash P60 (230-400 mesh) filled KP-Sil columns unless stated otherwise.

Concentration refers to the removal of volatile solvents via distillation using a rotary evaporator Büchi R-300HL equipped with a thermostated bath B-300, a vacuum regulator CVC-3000, followed by residual solvent removal under high vacuum.

$^{1}$H NMR (500 MHz) and $^{13}$C NMR (126 MHz) spectra were recorded at room temperature on a Bruker 500 MHz spectrometer. Data are represented as follows: chemical shift, multiplicity (s = singlet, d = doublet, t = triplet, q = quartet, m = multiplet, br = broad signal, bs = broad singlet, dd = doublet doublets, dt = doublet triplets, dq = doublet quartets, td = triplet doublets, ddd = doublet of doublet of doublets, ddt = doublet of doublet of triplets, dtd = doublet of triplet of doublets, dddd = doublet of doublet of doublet of doublets), coupling constants in Hertz (Hz). Chemical shifts (δ) are reported in ppm relative to the solvent peak. NMR spectra were analyzed using MestreNova© NMR data processing software (www.mestrelab.com).

High-resolution mass spectrometry (HR-MS) was performed by the analytical facility of the Chemistry department of the University of Basel on a Bruker maXis 4G QTOF ESI mass spectrometer.

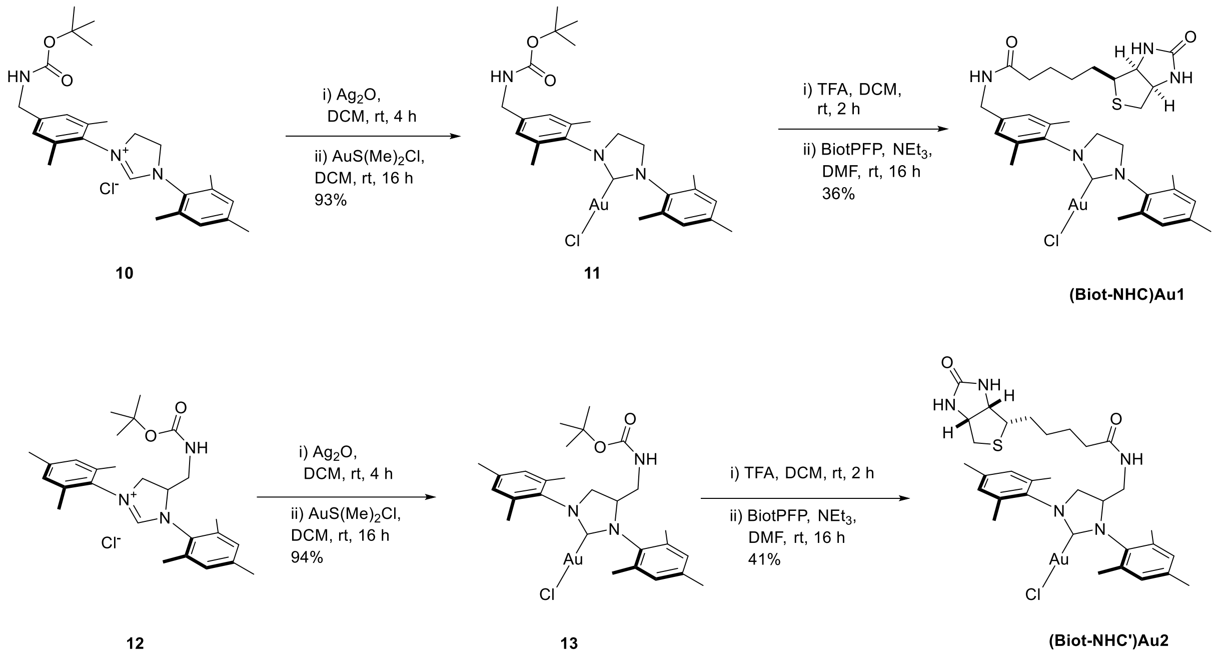


**Supplementary figure 10 | Synthesis routes for (Biot-NHC)Au1 and (Biot-NHC')Au2.**

NHC-salt **10** was prepared according to the procedure of Ward and co-workers in 6 steps[52]. NHC-salt **12** was prepared according to the procedure of Grubbs and co-workers in 4 steps[55]. An oven-dried reaction tube was charged with the corresponding NHC salt (1 eq), Silver(I)oxide (0.6 eq) and degassed DCM (10 mL/mmol). The mixture was stirred (4 h) at room temperature in the dark. The mixture was filtered through a plug of cellite into a tube containing chloro(dimethylsulfide) gold(I) (1 eq) and stirred in the dark (16 h). The mixture was filtered through a plug of cellite and the solvent was removed in vacuo to yield the corresponding gold complex as a white solid (**11**: 220 mg, 0.336 mmol, 93%; **13**: 126 mg, 0.189 mmol, 94%):

Boc-protected Gold complex **11**

$^1$H NMR (500 MHz, Methylene Chloride-d$_2$) δ 7.11 (s, 2H), 7.02 (s, 2H), 5.04 (s, 1H), 4.28 (d, J = 6.2 Hz, 2H), 4.02 (s, 4H), 2.37 (s, 6H), 2.34 (s, 6H), 2.33 (s, 3H), 1.46 (s, 9H).

$^{13}$C NMR (126 MHz, Methylene Chloride-d$_2$) δ 195.34, 140.97, 139.74, 136.94, 136.76, 136.28, 135.23, 130.19, 128.14, 51.32, 51.23, 28.68, 21.43, 18.47, 18.32.

HR-MS (ESI, pos): m/z calcd. for $C_{26}H_{35}AuClN_3O_2Na^+$ 676.1976 found 676.1970 $[M+Na^+]^+$.

Boc-protected Gold complex **13:**

$^1$H NMR (500 MHz, Methylene Chloride-d$_2$) δ 7.05 (s, 1H), 7.04 (s, 1H), 7.02 (s, 2H), 4.53 (m, 1H), 4.49 (m, 1H), 4.10 (t, J = 11.4 Hz, 1H), 3.94-3.81 (m, 1H), 3.47 (m, 1H), 3.26 (m, 1H), 2.41 (s, 3H), 2.36 (s, 4H), 2.34 (s, 4H), 2.33 (s, 3H), 2.32 (s, 4H), 2.31 (s, 3H), 1.36 (s, 9H).

$^{13}C$ NMR (126 MHz, Methylene Chloride-$d_2$) δ 196.01, 139.81, 136.78, 136.16, 130.75, 130.53, 130.23, 130.19, 63.81, 32.16, 28.47, 23.22, 21.43, 21.37, 19.53, 18.52, 18.38, 18.35, 14.44.

HR-MS (ESI, neg): m/z calcd. for $C_{27}H_{36}AuClN_3O_2^-$ 666.2156 found 666.2177 $[M-H^+]^-$; for $C_{27}H_{37}AuClN_3O_2HCOO^-$ 712.2211 found 712.2233 $[M+HCOO^-]^-$.

An oven-dried reaction tube was charged with the corresponding gold complex **11/13** (1 eq), degassed DCM (2 mL) and trifluoroacetic acid (20 eq). The mixture was stirred at room temperature in the dark until full deprotection. The solvent and the trifluoroacetic acid were fully removed via high vacuum. The dried crude was dissolved in degassed DMF (2 mL). Pentafluorobenzylbiotin (1 eq) and trimethylamine (20 eq) were added. The mixture was stirred (16 h) at room temperature in the dark. The solvent and the base were removed via high vacuum to yield a yellow crude. Purification via flash column chromatography (KP-Sil 10 g, gradient of 3% to 20% methanol in DCM) and subsequent precipitation of the product from DCM/hexane yielded the biotinylated gold complex as a white solid (**(Biot-NHC)Au1:** 67 mg, 0.086 mmol, 36%; **(Biot-NHC')Au2:** 18 mg, 0.023 mmol, 41%):

**(Biot-NHC)Au1:**

$^1H$ NMR (500 MHz, $CD_2Cl_2$) δ = 7.11 (s, 2H, C17H, C15H), 7.02 (s, 2H, C8H, C6H), 6.29 (s, 1H, N22H), 5.50 (s, 1H, N30H), 4.77 (s, 1H, N32H), 4.49 (dd, J = 8.0, 4.5 Hz, 1H, C33H), 4.38 (d, J = 6.0 Hz, 2H, C21H), 4.29 (dd, J = 7.9, 4.7 Hz, 1H, C29H), 4.01 (s, 4H, $C2H_2$, $C3H_2$), 3.17 (td, J = 7.5 Hz, 4.4, 2H, C28H), 2.92 (m, 1H, C34H), 2.68 (t, J = 12.2 Hz, 1H, C34H), 2.37 (s, 3H, $C20H_3$/$C19H_3$), 2.36 (s, 3H, $C19H_3$/$C20H_3$), 2.34 (s, 6H, $C10H_3$, $C12H_3$), 2.33 (s, 3H, $C11H_3$), 2.26 (td, J = 7.3, 2.2 Hz, 2H, $C24H_2$), 1.70-1.63 (m, 4H, $C25H_2$, $C27H_2$), 1.44 (m, 2H, $C26H_2$).

$^{13}C$ NMR (101 MHz, $CD_2Cl_2$) δ 195.01 (C1), 173.78 (C23), 164.60 (C31), 140.70 (C16), 139.68 (C4), 136.83 (C18, C14), 136.69 (C13), 136.27 (C5, C9), 135.24 (C7), 130.15 (C6, C8), 128.49 (C15/C17), 128.47 (C17/C15), 62.21 (C29), 60.81 (C33), 56.13 (C28), 51.32 (C3/C2), 51.27 (C2/C3), 43.15 (C21), 41.11 (C34), 36.40 (C24), 28.70 (C25, C26, C27), 21.42 (C11), 18.51 (C20/C19), 18.47 (C19/C20), 18.35 (C10, C12).

HR-MS (ESI, pos): m/z calcd. for $C_{31}H_{41}AuN_5O_2S^+$ 744.2641 found 744.2651 $[M-Cl^-]^+$.

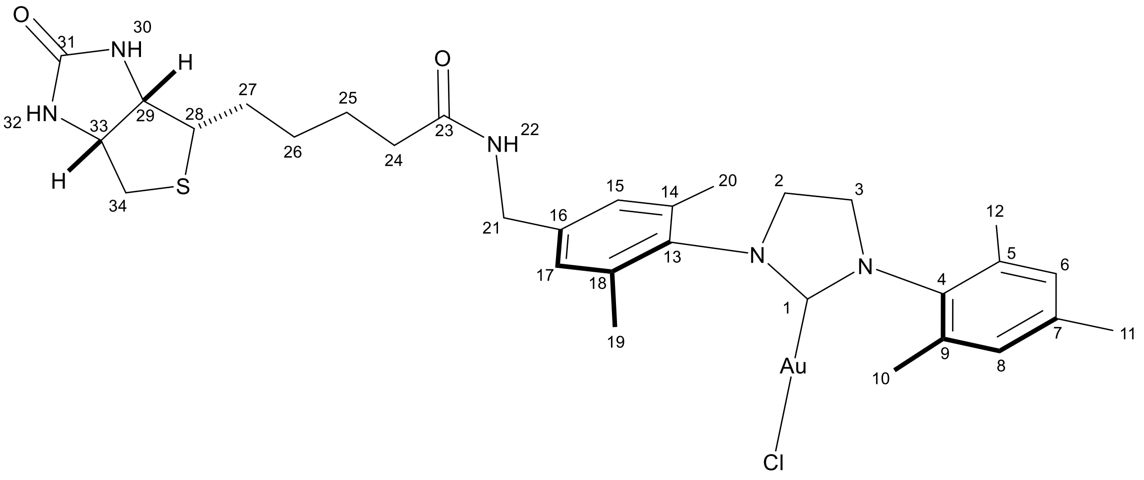


**Supplementary figure 11 | Structure of (Biot-NHC)Au1.**

**(Biot-NHC')Au2**:

$^{1}$H NMR (500 MHz, Methylene Chloride-$d_2$) δ 7.06 (s, 1H, C8H/C6H), 7.04 (s, 1H, C6H/C8H), 7.01 (s, 2H, C15H, C17H), 5.71 (d, J = 6.1 Hz, 1H, N23H), 5.38 (s, 1H, N31H), 4.76 (d, J = 5.3 Hz, 1H, N33H), 4.53 (m, 1H, C3H), 4.45 (m, 1H, C34H), 4.24 (m, 1H, C30H), 4.11 (td, J = 11.3, 4.4 Hz, 1H, C2H), 3.82 (ddd, J = 11.6, 8.6, 7.7 Hz, 1H, C2H), 3.74 (m, 1H, C22H), 3.24 (m, 1H, C22H), 3.12 (m, 1H, C29H), 2.89 (dt, J = 12.9, 5.0 Hz, 1H, C35H), 2.65 (dd, J = 12.8, 7.5 Hz, 1H, C35H), 2.43 (s, 3H, $C10H_3/C11H_3$), 2.36 (s, 3H, $C19H_3/C20H_3$), 2.35 (s, 3H, $C12H_3$), 2.33 (s, 3H, $C21H_3$), 2.31 (s, 6H, $C11H_3/C10H_3$, $C20H_3/C19H_3$), 2.01 (m, 2H, $C25H_2$), 1.61 (s, 2H, $C28H_2$), 1.47 (s, 2H, $C26H_2$), 1.33 (s, 2H, $C27H_2$).

$^{13}$C NMR (126 MHz, Methylene Chloride-$d_2$) δ 196.04 (C1), 173.90 (C24), 163.61 (C32), 139.80 (C7, C16), 136.82 (C4), 136.36 (C13), 135.13 (C14, C18), 134.41 (C9, C5), 130.76 (C8/C6), 130.53 (C6/C8), 130.16 (C15, C17), 63.40 (C3), 62.25 (C30), 60.60 (C34), 55.91 (C29), 55.36 (C2), 42.47 (C22), 41.15 (C35), 36.00 (C25), 28.58 (C27/C28), 28.51 (C28/C27), 25.68 (C26), 21.43 (C12, C21), 19.59 (C10/C11), 18.53 (C19/C20), 18.38 (C11/C10), 18.34 (C20/C19).

HR-MS (ESI, pos): m/z calcd. for $C_{32}H_{43}AuN_5O_2S^+$ 758.2803 found 758.2794 $[M\text{-}Cl^-]^+$; for $C_{32}H_{43}AuClN_5O_2SNa^+$ 816.2384 found 816.2384 $[M+Na^+]^+$.

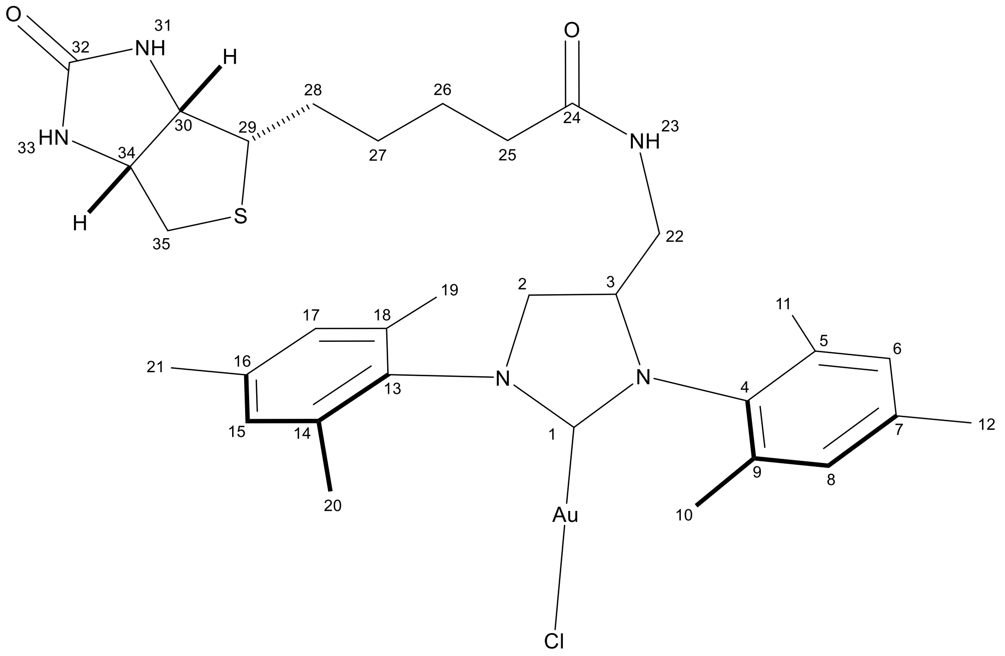

**Supplementary figure 12 | Structure of (Biot-NHC')Au2.**

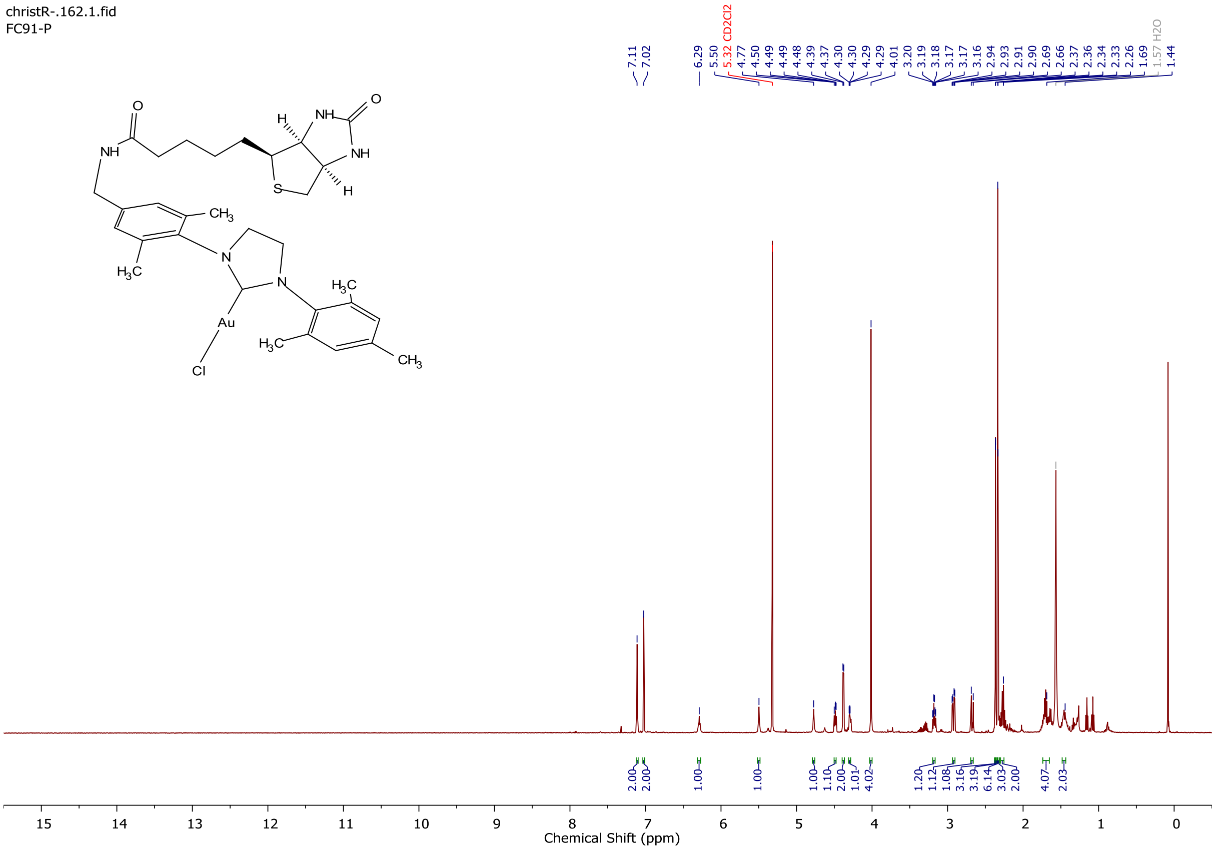


**Supplementary figure 13 | $^{1}$H NMR spectrum of (Biot-NHC)Au1 in methylene chloride-d$_2$.**

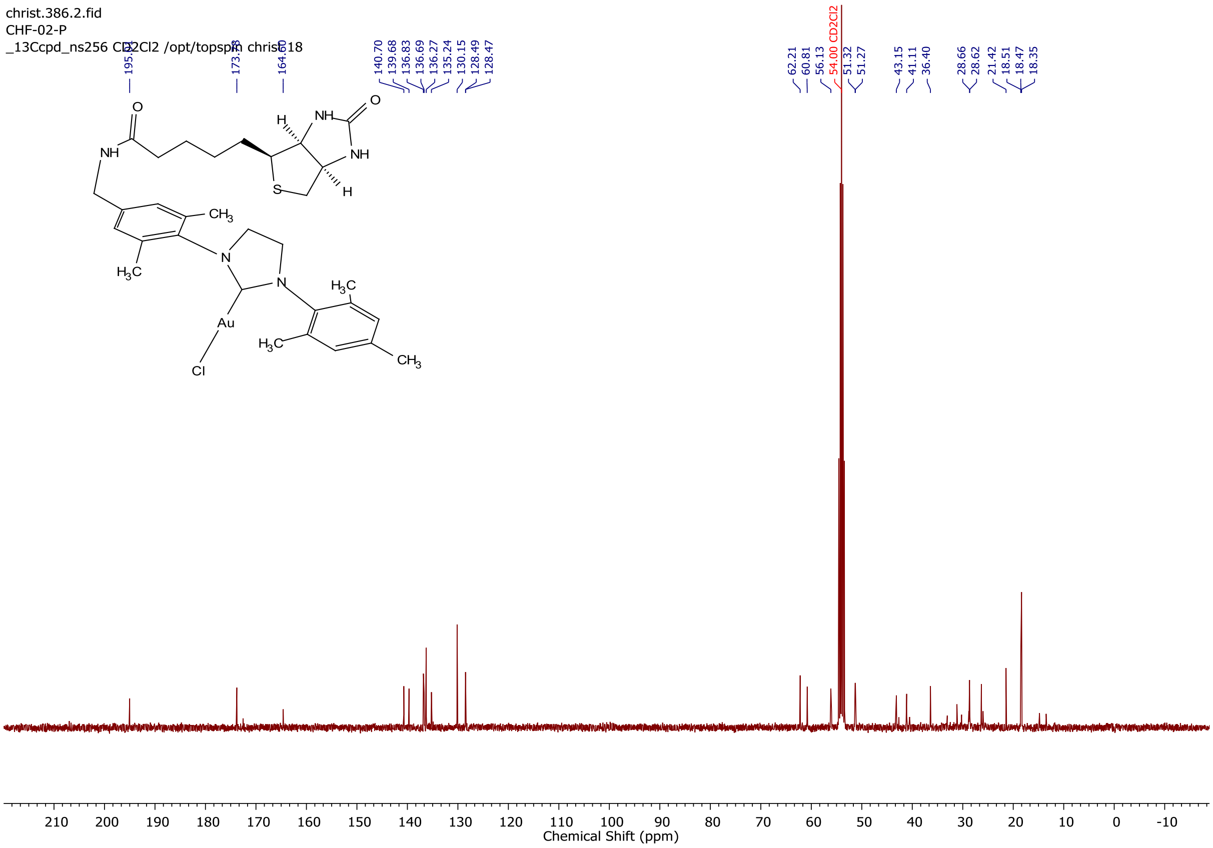


**Supplementary figure 14 | $^{13}$C NMR spectrum of (Biot-NHC)Au1 in methylene chloride-d2.**

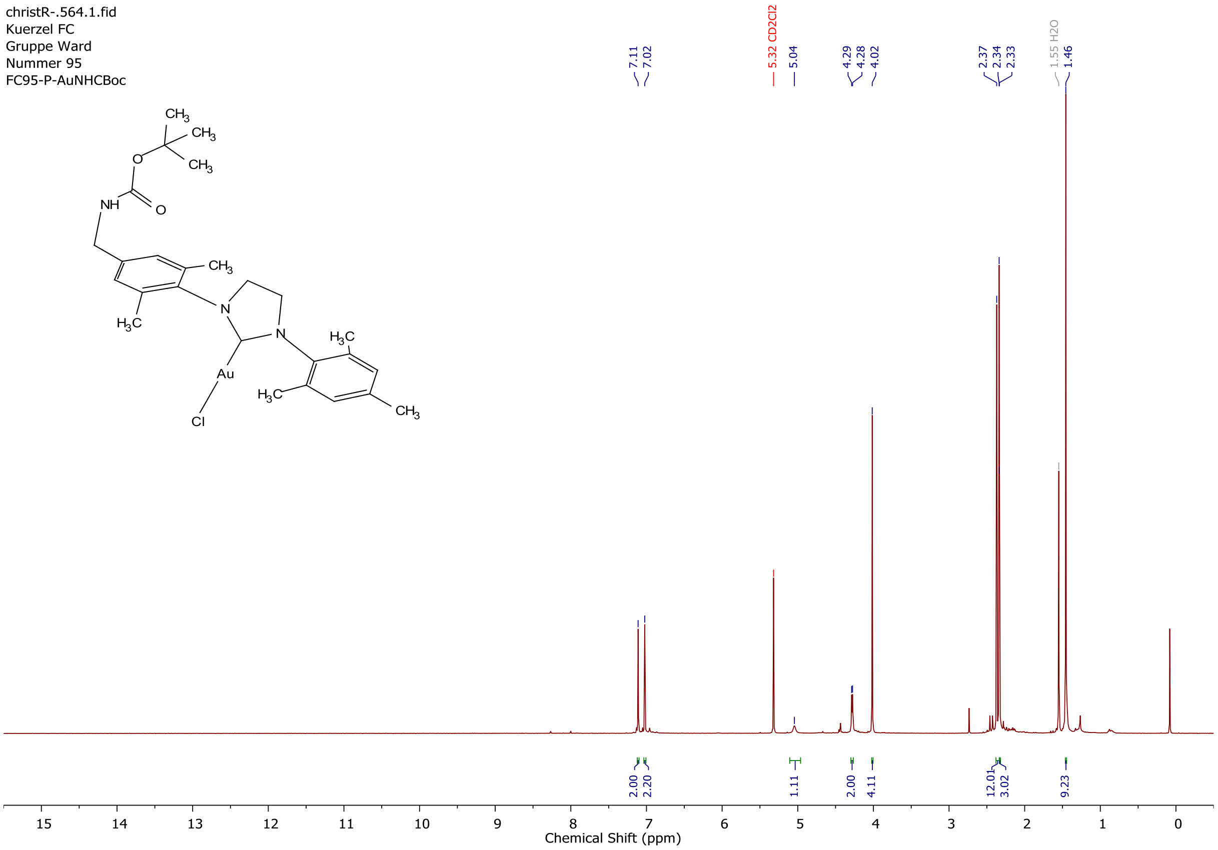


**Supplementary figure 15 | ¹H NMR spectrum of gold complex 11 in methylene chloride-d2.**

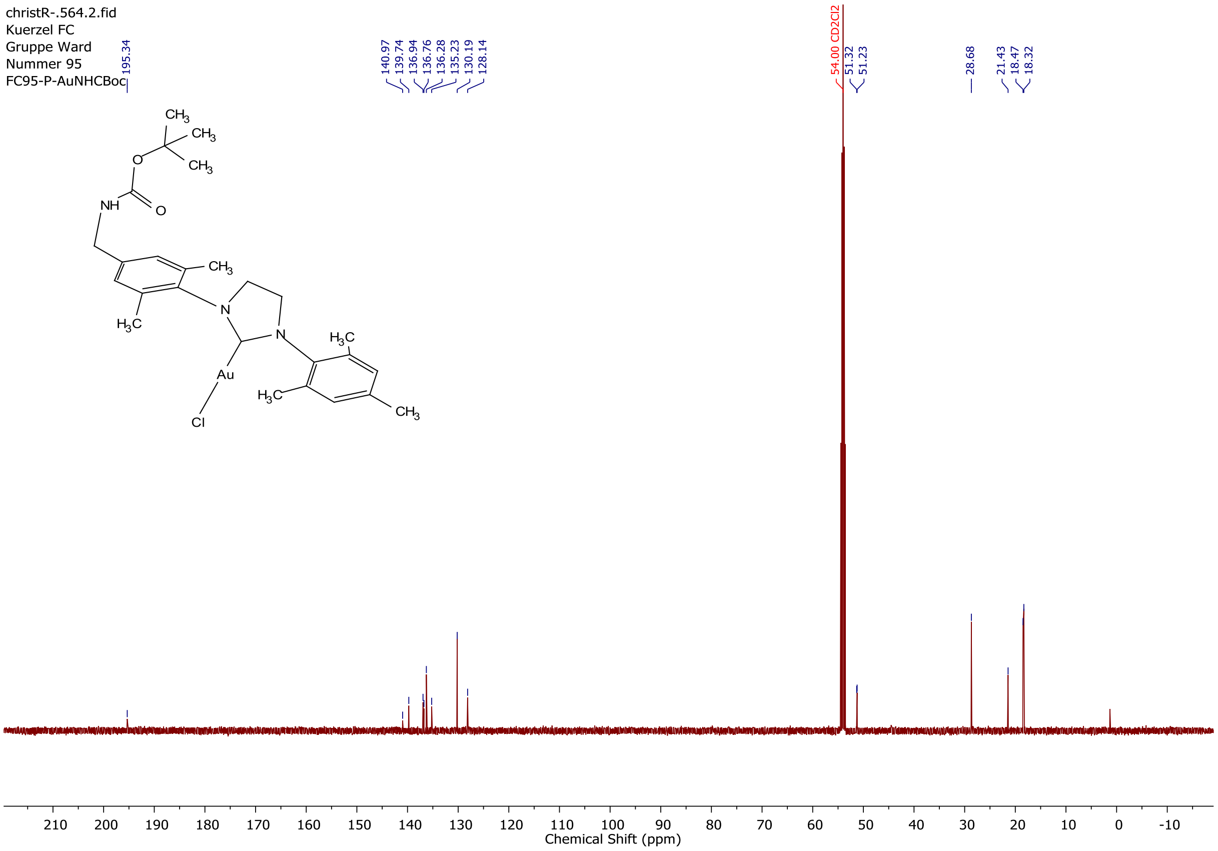


**Supplementary figure 16 | ¹³C NMR spectrum of gold complex 11 in methylene chloride-d2.**

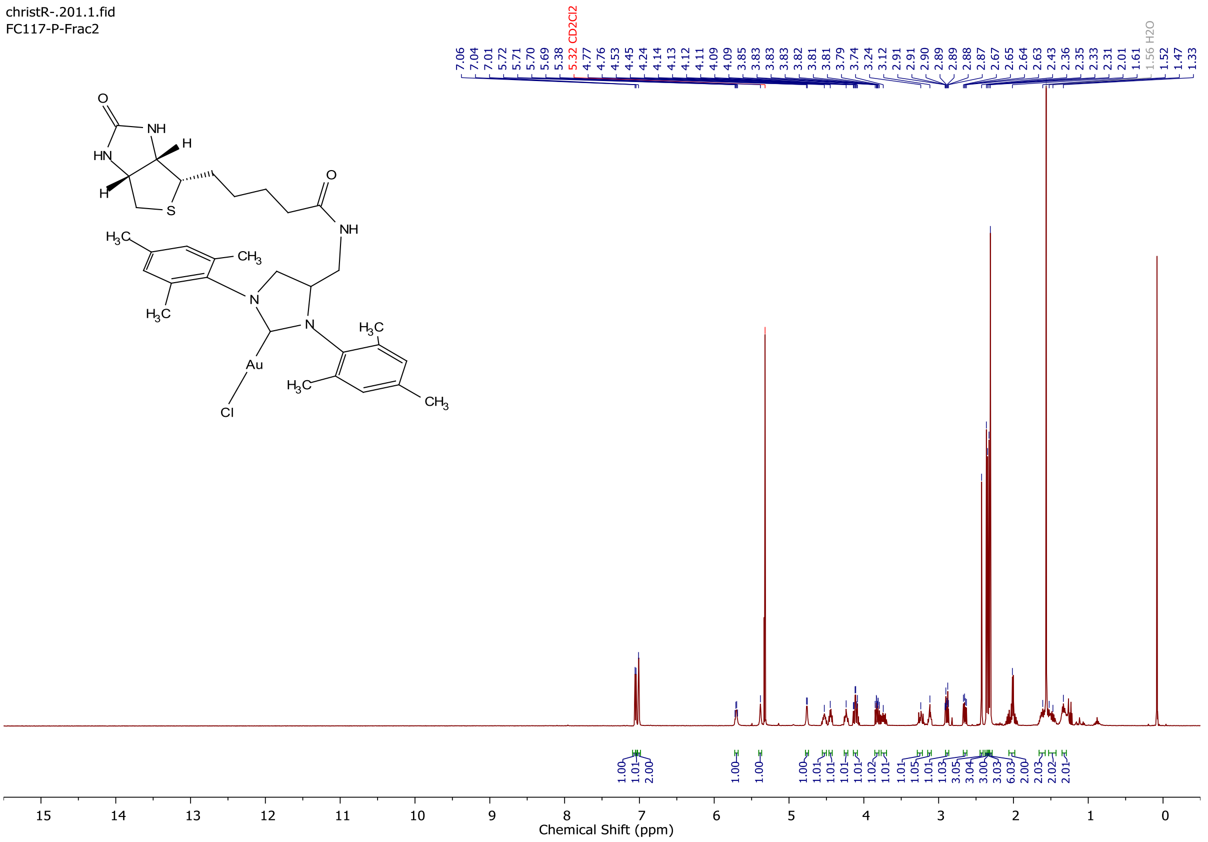


**Supplementary figure 17 | $^{1}$H NMR spectrum of (Biot-NHC')Au2 in methylene chloride-d2.**

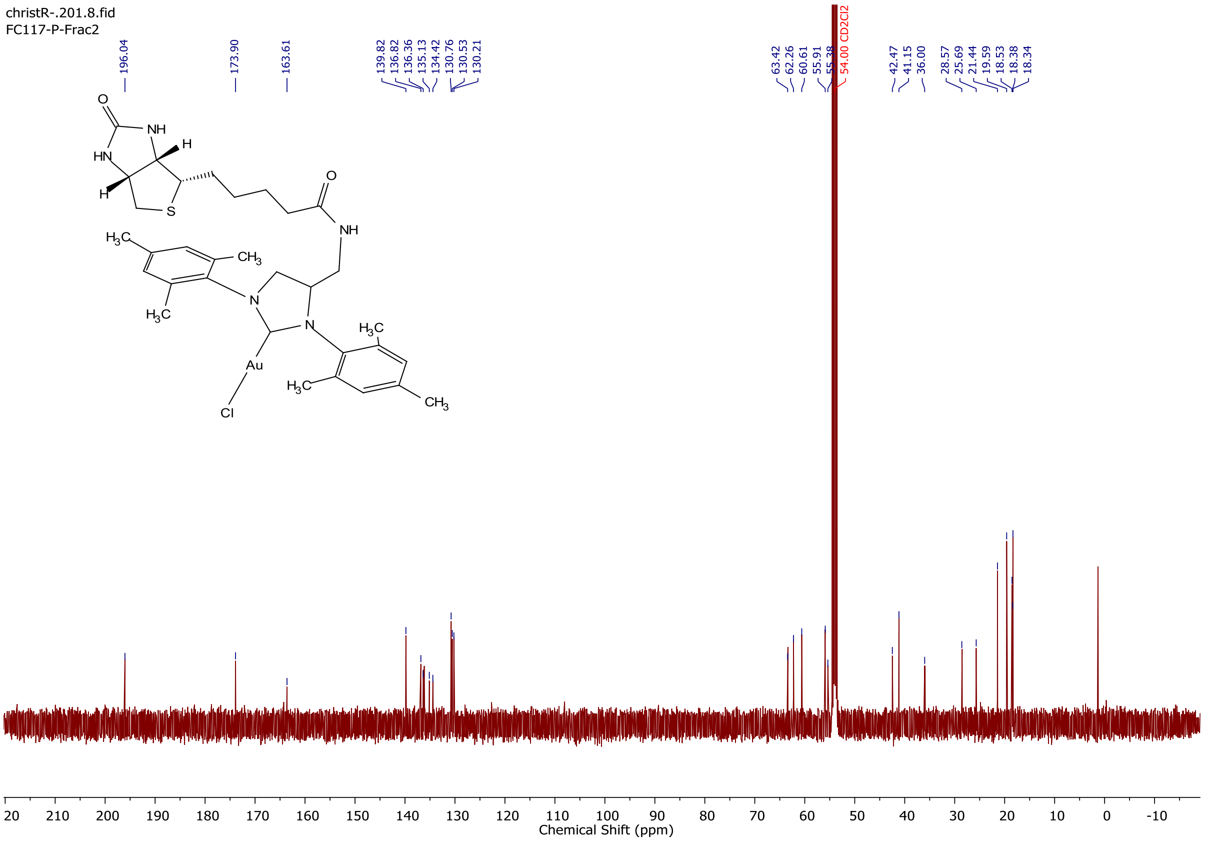


**Supplementary figure 18 | $^{13}$C NMR spectrum of (Biot-NHC')Au2 in methylene chloride-d2.**

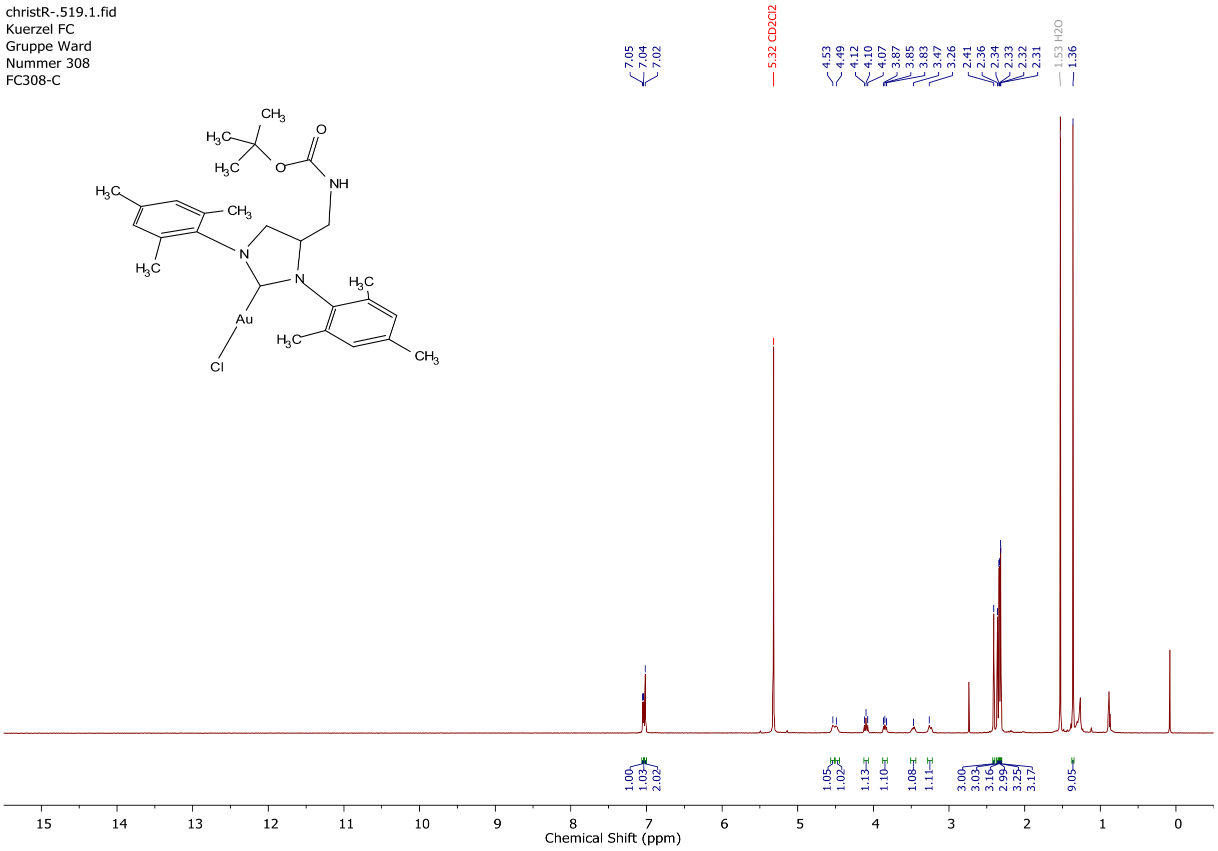


**Supplementary figure 19 | $^{1}$H NMR spectrum of gold complex 13 in methylene chloride-d2.**

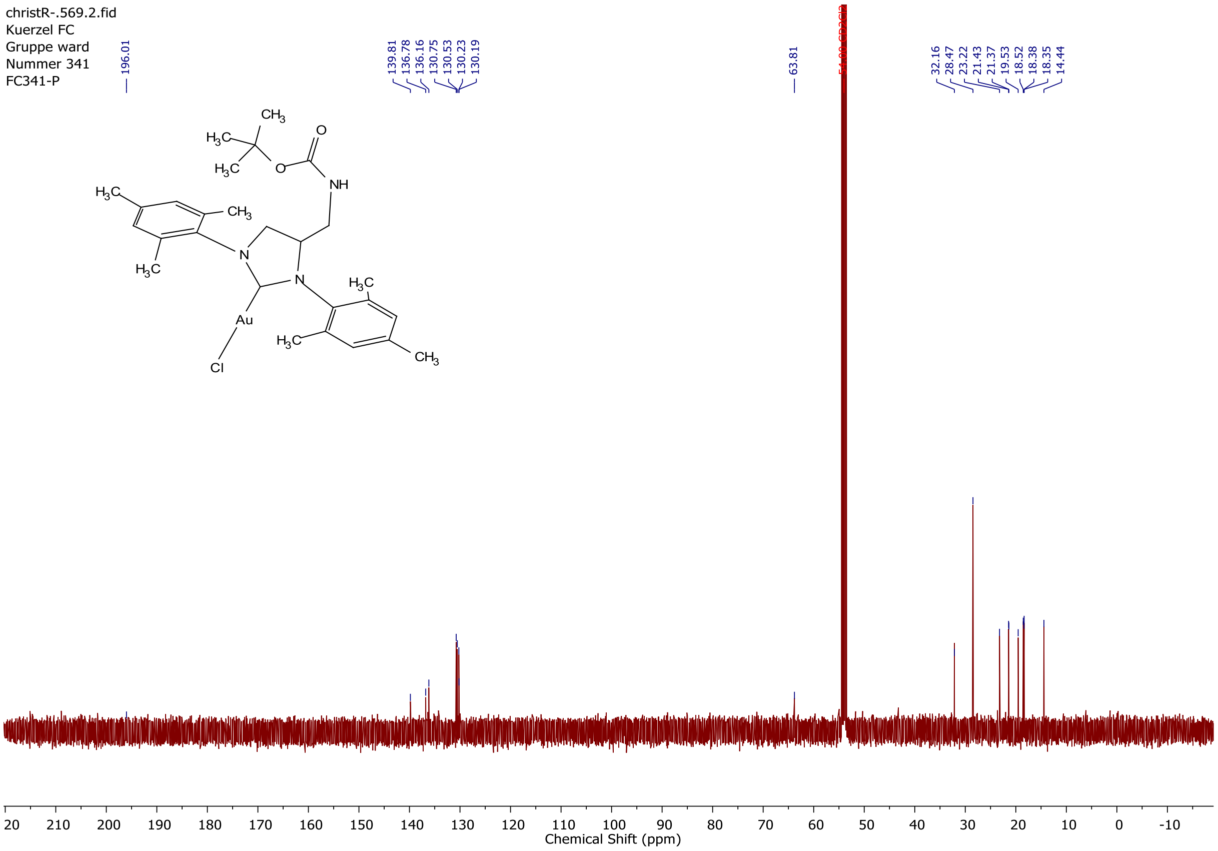


**Supplementary figure 20 | $^{13}$C NMR spectrum of gold complex 13 in methylene chloride-d2.**

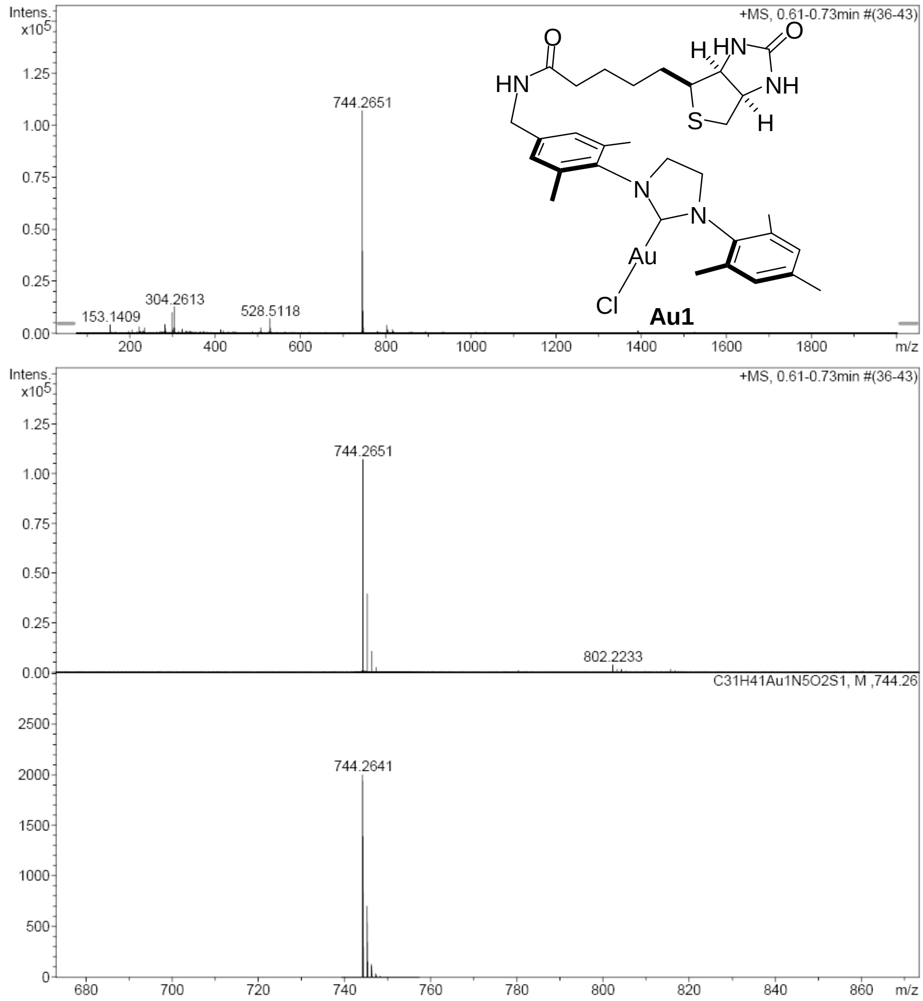


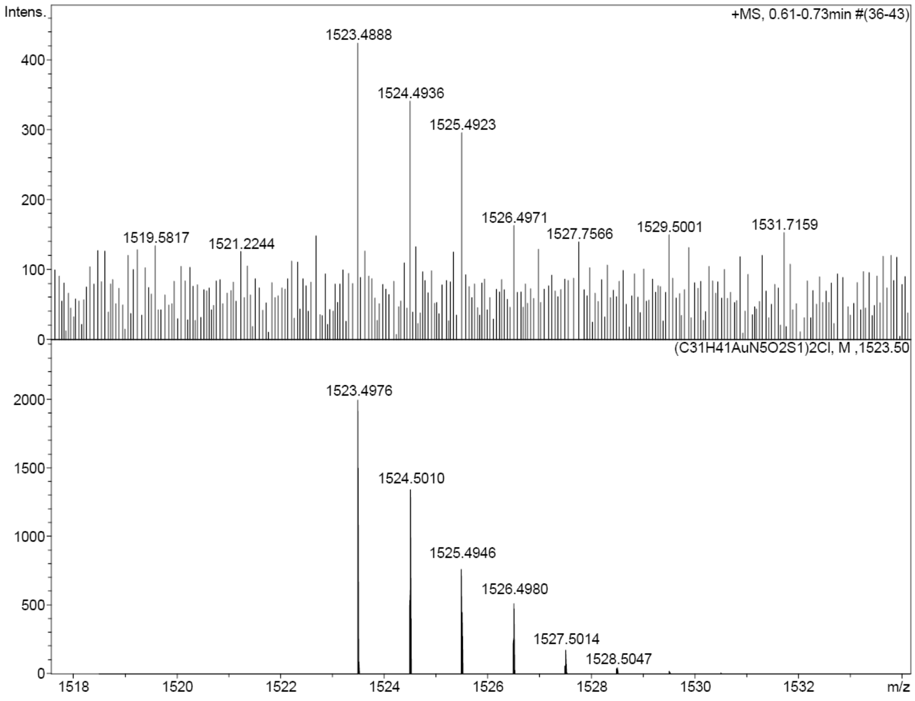


**Supplementary figure 21 | HR-MS spectrum of (Biot-NHC)Au1.**

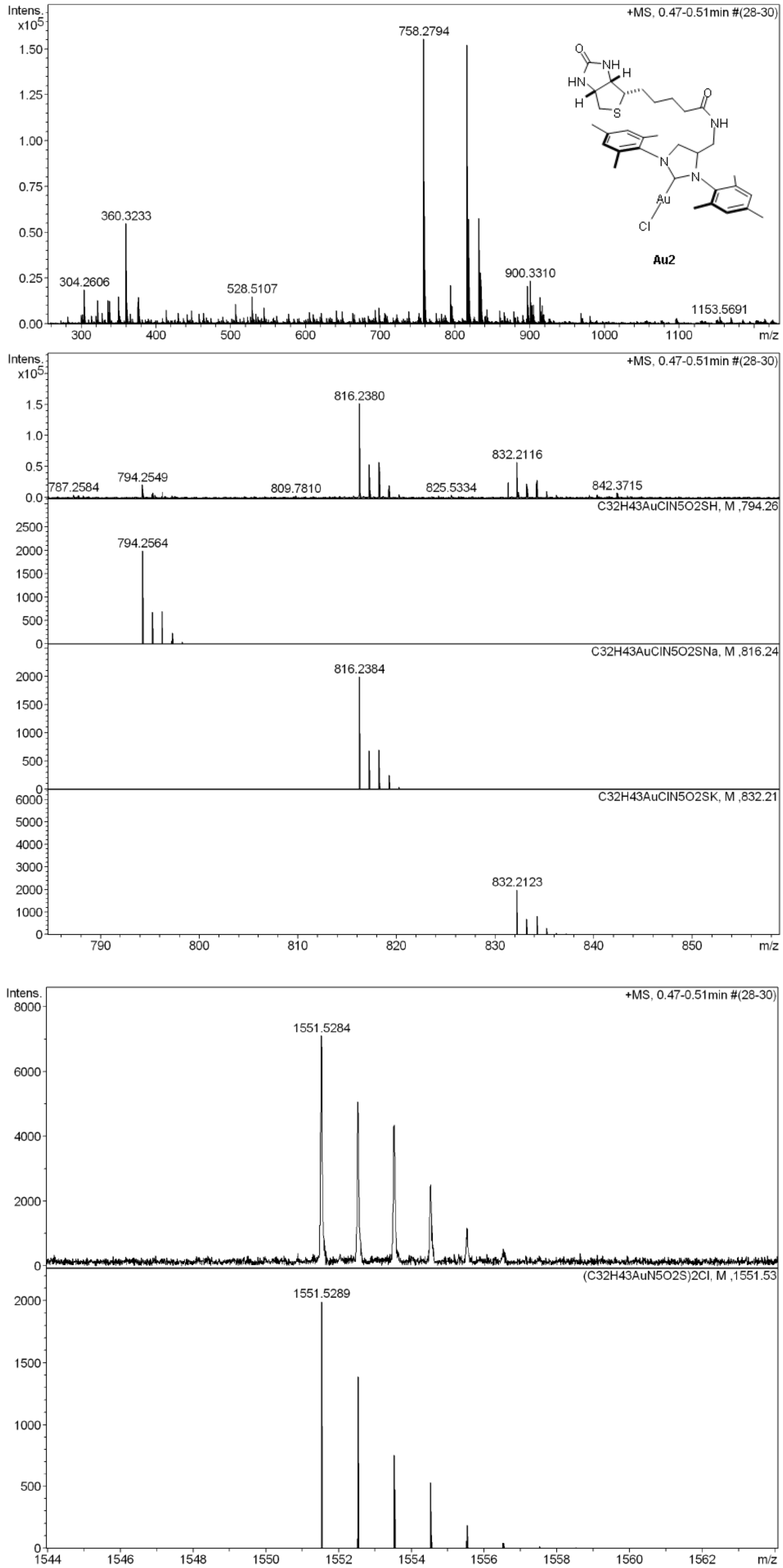


**Supplementary figure 22 | HR-MS spectrum of (Biot-NHC')Au2.**

#### Synthesis of substrate 8 and product 9

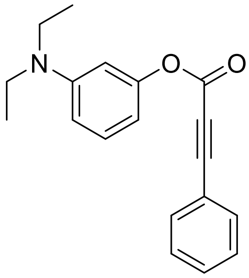


**3-(diethylamino)phenyl 3-phenylpropiolate (8):** Prepared according to a modified procedure of Do et al[56]. To a solution of 3-diethylaminophenol (826 mg, 5.00 mmol, 1.0 eq) in dry DCM (20 mL) was added a solution of 3-phenylpropiolic acid (804 mg, 5.50 mmol, 1.1 eq) in dry DCM (10 mL). The solution was cooled to 0 °C before the dropwise addition of a solution of DMAP (305 mg, 2.50 mmol, 0.5 eq) and DCC (1.55 g, 7.50 mmol, 1.5 eq) in dry DCM (10 mL). The solution was stirred at room temperature for 16 h, then cooled to 0 °C and filtered to remove most of the formed dicyclohexylurea. The crude mixture was concentrated in vacuo and purified via automated flash column chromatography (KP-Sil50g) to yield ester **8** as an orange oil (622 mg, 2.12 mmol, 42%).

$^1$H NMR (500 MHz, DMSO-$d_6$) δ 7.71 – 7.66 (m, 2H), 7.63 – 7.58 (m, 1H), 7.52 (t, J = 7.7 Hz, 2H), 7.19 (t, J = 8.2 Hz, 1H), 6.58 (dd, J = 8.4, 2.5 Hz, 1H), 6.50 (t, J = 2.3 Hz, 1H), 6.40 (dd, J = 7.8, 2.1 Hz, 1H), 3.32 (q, J = 7.2 Hz, 4H), 1.07 (t, J = 7.0 Hz, 6H).

$^{13}$C NMR (126 MHz, DMSO-$d_6$) δ 151.74, 151.17, 148.70, 132.95, 131.62, 129.96, 129.17, 118.15, 109.43, 107.36, 104.27, 87.95, 80.26, 43.73, 12.27.

HR-MS (ESI, pos): m/z calcd. for $C_{19}H_{19}NO_2H^+$ 294.1489 found 294.1487 $[M+H^+]^+$; m/z calcd. for $C_{19}H_{19}NO_2Na^+$ 316.1308 found 316.1306 $[M+Na^+]^+$.

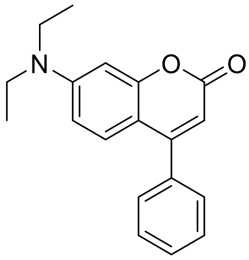


**7-(diethylamino)-4-phenyl-2H-chromen-2-one (9):** Prepared according to a modified procedure of Do et al[56]. In a reaction tube, ester **8** (96.8 mg, 0.33 mmol, 1.0 eq) was diluted with dry DCM (1.6 mL) and dry EtOH (0.4 mL) before adding hydrogen tetrachloroaurate(III) hydrate (23.6 mg, 0.07 mmol, 0.2 eq). The reaction mixture was stirred at room temperature for 16 h in the dark. The solvents were removed in vacuo and the crude was purified via automated flash column chromatography (KP-Sil25g) to yield coumarin **9** as a yellow oil (43 mg, 0.15 mmol, 44%).

$^{1}H$ NMR (500 MHz, DMSO-$d_6$) δ 7.54 (dt, J = 5.0, 2.2 Hz, 3H), 7.51 – 7.46 (m, 2H), 7.18 (d, J = 9.1 Hz, 1H), 6.68 (dd, J = 9.1, 2.6 Hz, 1H), 6.61 (d, J = 2.5 Hz, 1H), 5.93 (s, 1H), 3.43 (q, J = 7.0 Hz, 4H), 1.12 (t, J = 7.0 Hz, 6H).

$^{13}C$ NMR (126 MHz, DMSO-$d_6$) δ 160.62, 156.37, 155.59, 150.50, 135.61, 129.42, 128.77, 128.30, 127.67, 108.91, 107.26, 106.77, 97.13, 44.02, 12.32.

HR-MS (ESI, pos): m/z calcd. for $C_{19}H_{19}NO_2H^+$ 294.1489 found 294.1491 $[M+H^+]^+$; m/z calcd. for $C_{19}H_{19}NO_2Na^+$ 316.1308 found 316.1310 $[M+Na^+]^+$.

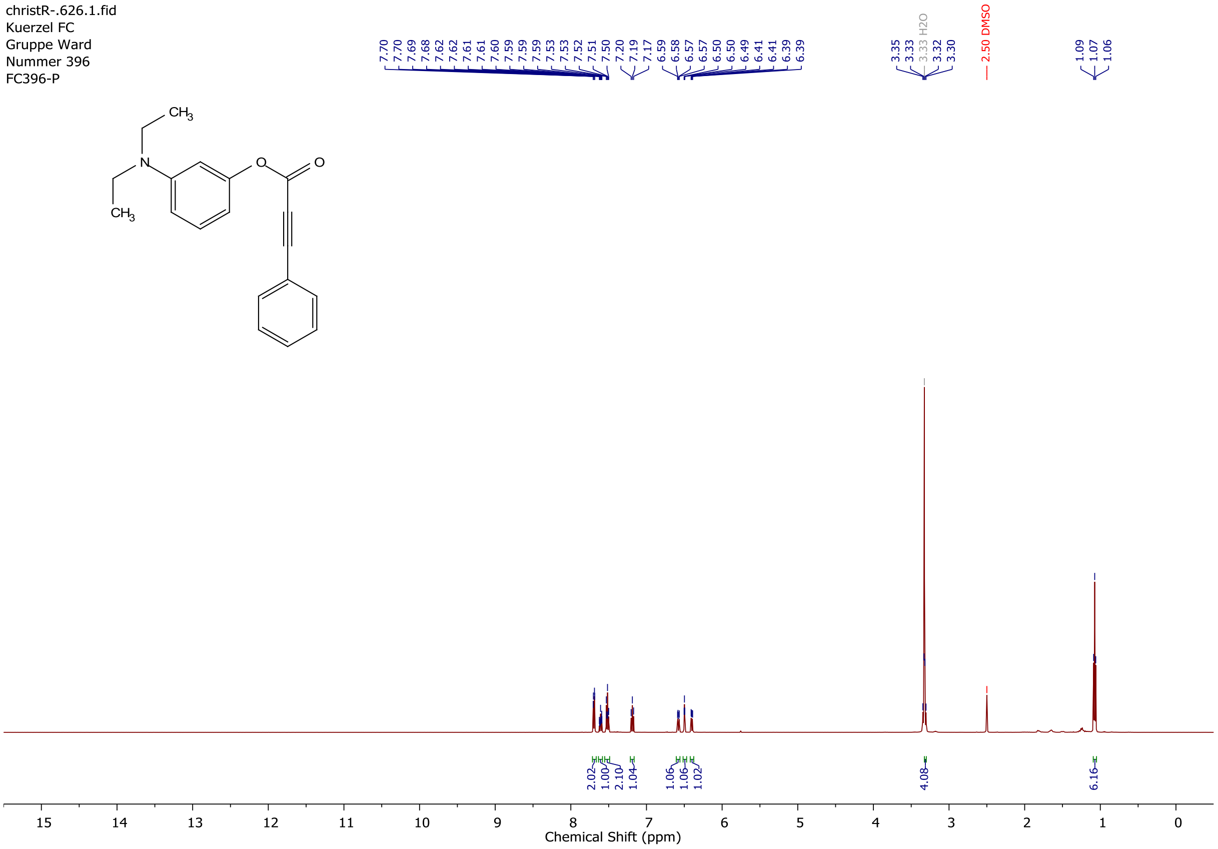


**Supplementary figure 23 | H-NMR spectrum of ester 8 in DMSO-d6.**

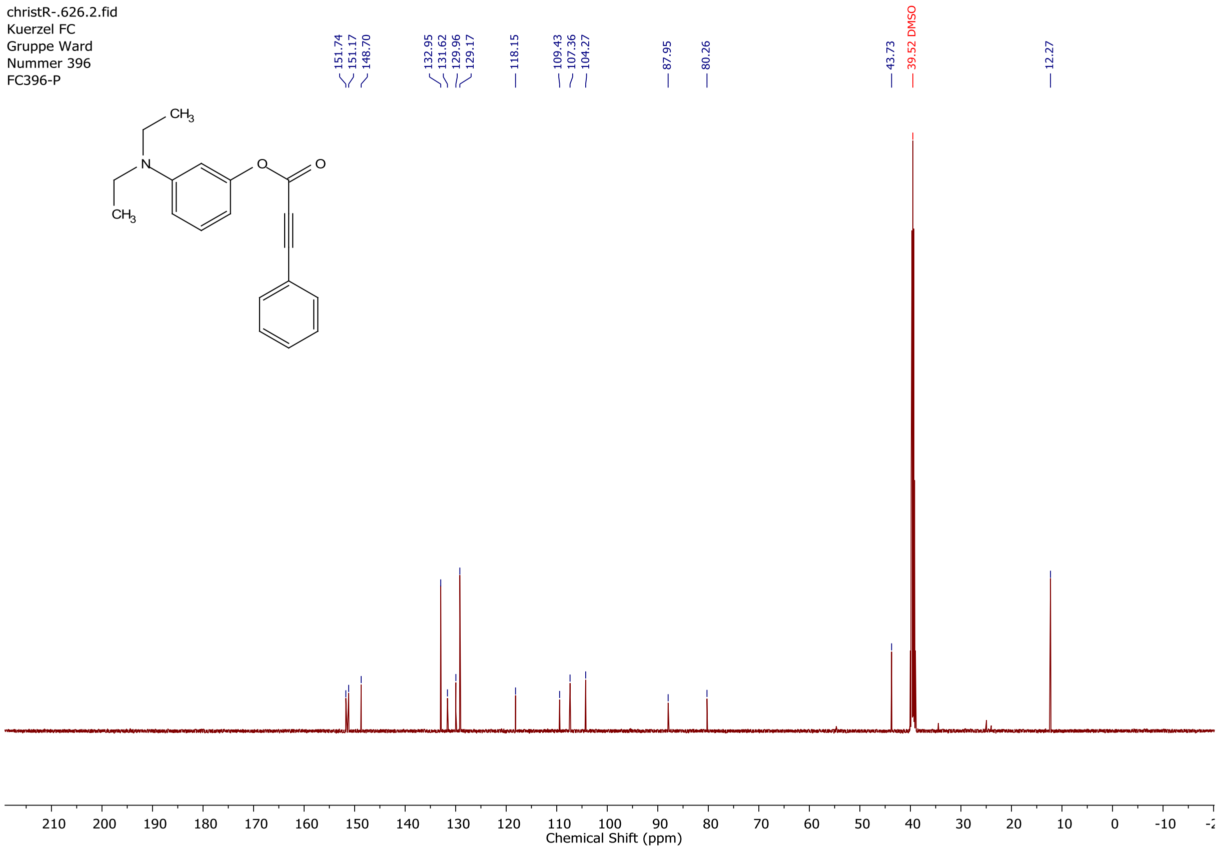


**Supplementary figure 24 | C-NMR spectrum of ester 8 in DMSO-d6.**

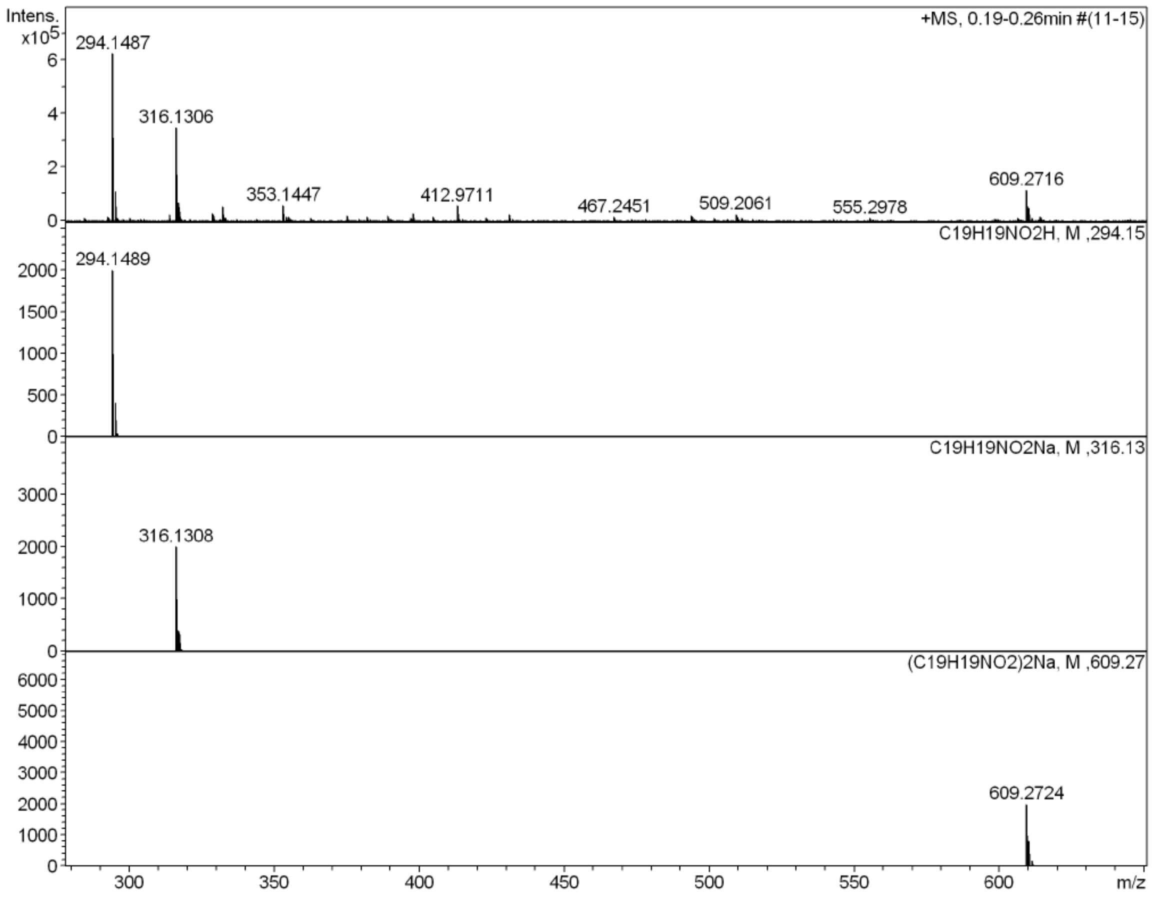


**Supplementary figure 25 | HR-MS spectrum of ester 8.**

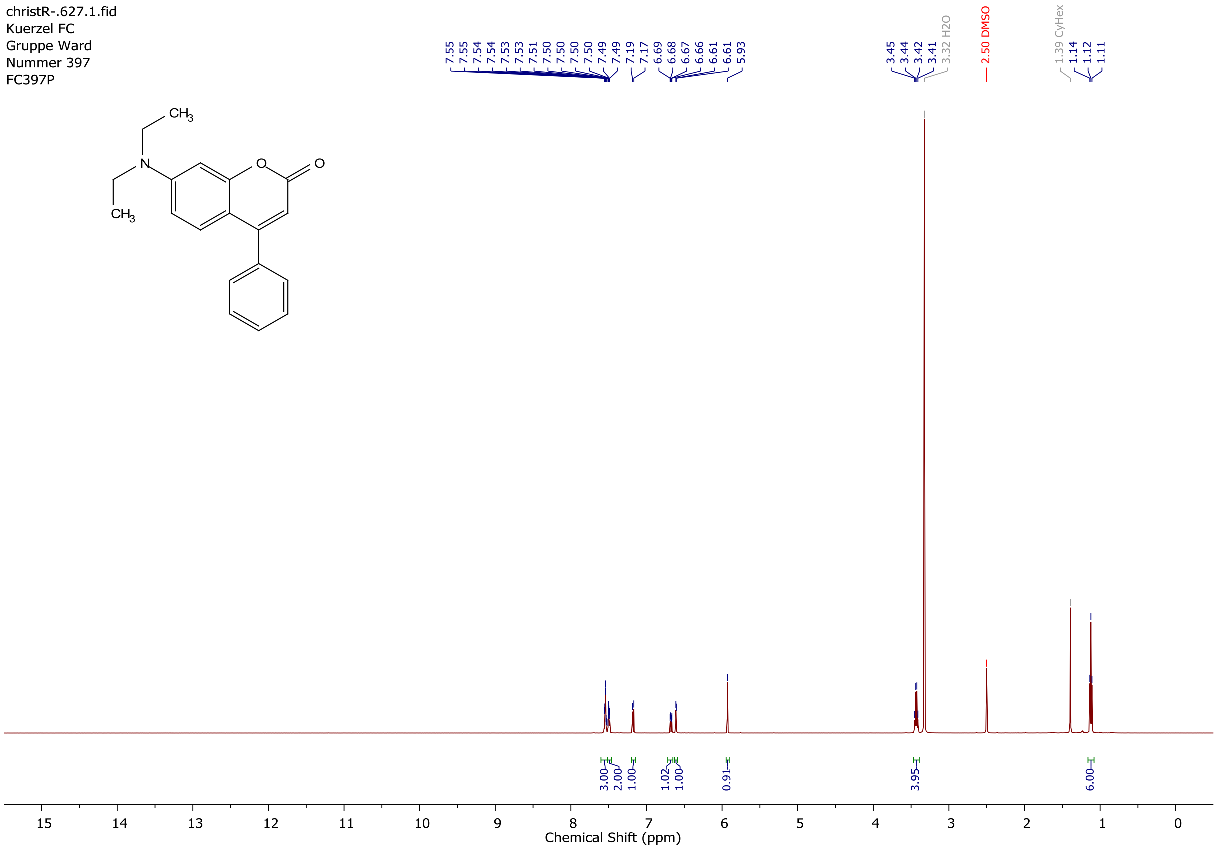


**Supplementary figure 26 | H-NMR spectrum of coumarin 9 in DMSO-d6.**

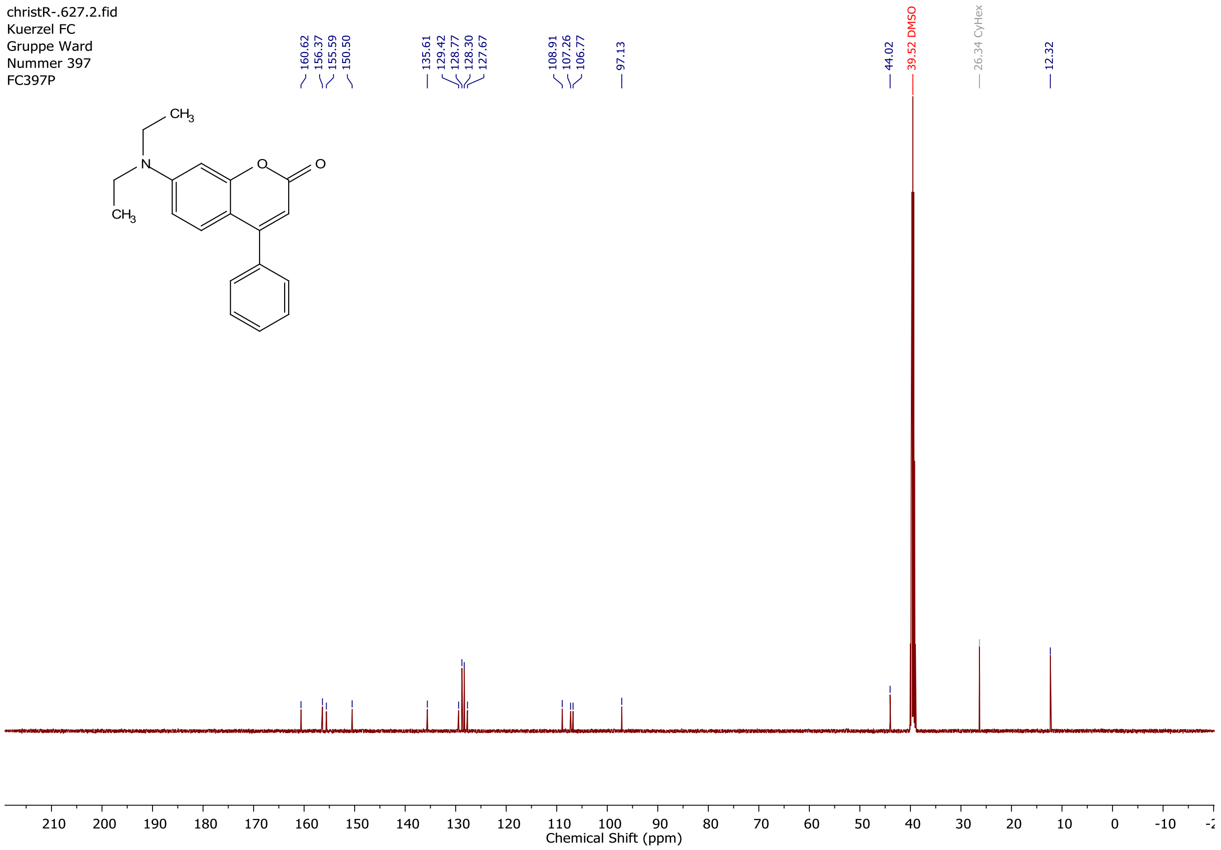


**Supplementary figure 27 | C-NMR spectrum of coumarin 9 in DMSO-d6.**

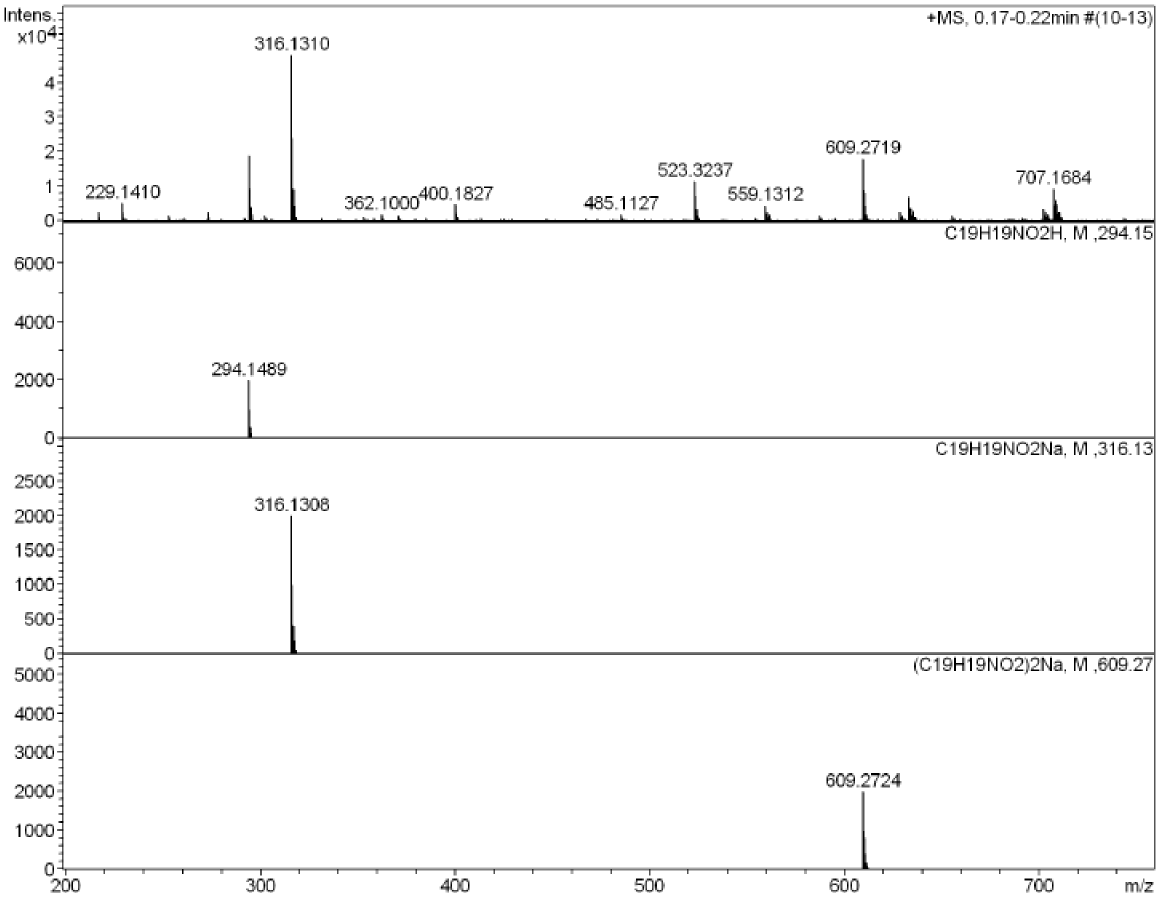


**Supplementary figure 28 | HR-MS spectrum of coumarin 9.**

#### Supplementary References


55. Jordan, J. P. & Grubbs, R. H. Small-molecule N-heterocyclic-carbene-containing olefin-metathesis catalysts for use in water. *Angew. Chem. Int. Ed.* **46**, 5152–5155 (2007).

56. Do, J. H., Kim, H. N., Yoon, J., Kim, J. S. & Kim, H. J. A rationally designed fluorescence turn-on probe for the gold(III) ion. *Org. Lett.* **12**, 932–934 (2010).
